# Getting there and staying there: supporting and enabling persistent human life on Mars using synthetic natural rubber, self-healing materials, and biological batteries

**DOI:** 10.1101/345496

**Authors:** Nischal Acharya, Natalie Baker, Marilu Krystal Bravo, Katie Gu, Sierra Harken, Michael Howland Herschl, Addie Petersen, Ileana Pirozzi, Dylan Spangle, Gordon Sun, Brian Vuong, Nils J.H. Averesch, Kosuke Fujishima, Trevor! J. Kalkus, Kara J. Helmke Rogers, Lynn J. Rothschild

## Abstract

Planetary exploration requires a balance between preemptive planning and financial feasibility. The risk of mid-mission equipment failure, power shortages, or supply depletion incentivizes precautionary measures, but the financial strain of sending unnecessary mass into space limits this practice.

To balance the two, our team explored the advantages of biological solutions, namely the self-sustaining abilities of low-mass organisms, to make planetary exploration more self-sufficient and economical. Prioritizing repair over replacement, we are developing self-healing materials embedded with *Bacillus subtilis*. For longer-lasting energy, we are designing a “biobactery” using linearly oriented *Escherichia coli* to generate power. For renewable materials, we are engineering bacteria to synthesize and degrade rubber. Individually, these projects offer sustainable alternatives for repair, power, and materials. But when combined, these consolidated insights can provide us with the power to get to Mars and resources to sustain us while we’re there.

## Introduction

### BioRubber Intro

Over 25 million tons of rubber are produced globally per annum, and demand is rising. Close to half of that rubber is harvested from *Hevea brasiliensis* (“natural rubber”), with the remaining rubber derived from petroleum-based sources (deemed “synthetic rubber”) (Beilen et al. 2007). To meet demand, more than 1.23 million hectares of land have been converted to *Hevea* plantations, mostly in Southeast Asia where human rights violations have been identified by the UN High Commissioner for Refugees. Demand is so high that *Hevea* land usage is estimated to double by 2050, threatening biodiversity and potentially compromising water resources, according to a report in *Science*. (Ziegler 2009) Synthetic rubber does not fare much better, producing toxic wastes and dependency on foreign oil.

Without any rubber trees or petroleum as raw material sources, rubber production in space is far more difficult, but may be necessary for extended habitation. Rubber has myriad uses in a space exploration/colonization setting: incorporation into hoses, sealing products, construction, footwear, aerospace technology, scientific equipment. Only a limited amount of spare material can be carried by each mission, and space and mass are both at a premium. Therefore there is great interest in creating a new method of rubber production, independent of Earth’s natural resources which cannot be reproduced in a lab.

Commercial interest in synthetic biology in the rubber industry boomed the past decade due to the volatility of petroleum and *Havea* markets, as well as increased emphasis on corporate social responsibility and climate goals. Partnerships between Genencor and Dupont, as well as Goodyear and Amyris have successfully produced rubber monomers (i.e. isoprene). Unlike our team, however, these companies assemble the rubber using chemical catalysts. This commercial exploration of a synthetic biology solution by major players in the rubber industry demonstrates the potential for commercial applications of our product.

Additional work is also being conducted both in academia and at the US government level exploring methods of rubber production synthetically, and from other organisms (notably in *Taraxacum kok-saghyz* and *Parthenium argentatum*), and characterizing rubber produced from these alternative sources.

Several iGEM teams have made progress towards synthetic biological rubber production, as well as methods for recycling rubber for greater sustainable use. The Denmark 2013 iGEM team developed constructs which allowed for direct production of rubber in *E. coli*. In 2015, Brasil-USP focused on industry practices, undertaking “RubberBye”, which focused on the degradation of natural rubber. Then, in 2016, Stanford-Brown further sought to expand and refine the work of Denmark 2013. They added a second *Hevea* gene in a new construct, as well as a secondary construct to significantly increase the quantity of rubber production.

Building upon the successes of previous iGEM teams and industry, our vision was to produce a rubber product that can be *created* and *degraded* using synthetic biology. In developing this product, we set out to avoid the limitations found on earth, and grant an opportunity to make rubber more sustainable for space exploration.

One of our main focuses was eliminating the need for a cell lysis step, something which required impractical reagents, and added unnecessary mass for missions in space. We sought to do this with two steps. The first, based upon information provided to us by an industry-member we interviewed, was to act on the assumption that our cells could secrete our isoprene monomers: did this by having our cells secreting the isoprene monomers: isopentenyl-pyrophosphate (IPP) and dimethylallyl-pyrophosphate (DMAPP). Knowing this, we then tried to engineer the protein responsible for polymerization of these monomers - cis-prenyltransferase - to be expressed on the cell’s outer membrane, fused to the outer membrane protein ompA, which would effectively move our cellular machinery to the outside of our *E. coli.* This was inspired by Su et al., 2013, in which high expression of the human receptor activity-modifying protein RAMP1 was expressed on the outside of *E. coli* as a fusion with the similar outer membrane protein ompF.

In acting upon our design, we created a construct that would express a protein fusion of ompA and components of the composite construct (BBa_K2027039) that had been developed by our predecessor team, 2016 Stanford-Brown iGEM Team. The composite construct from 2016 (referred to as Latex Operon below) contained three cDNA sequences originally found in the rubber plant, *Hevea brasiliensis*, that encoded for small rubber particle protein (SRPP), *Hevea* rubber transferase 1 (HRT1), and *Hevea* rubber transferase 2 (HRT2), respectively.

HRT1 and HRT2 are components of the enzyme *cis*-prenyltransferase, which is involved in the polymerization of *cis*-1,4-polyisoprene from the isoprene monomers isopentenyl-pyrophosphate (IPP) and dimethylallyl-pyrophosphate (DMAPP). Previous research indicates that both HRT1 and HRT2 contribute to natural rubber formation despite there being possible differences in the functional efficiency and mechanisms of the two rubber transferases. One study on laticifer-specific gene expression analyzed various tissues as well as latex from rubber-producing plants. In this study, HRT1 transcripts were only found in the latex of tissues, whereas HRT2 transcripts were found in all the tissues analyzed. Despite there being a high degree of nucleotide similarity between HRT1 and HRT2, these results suggest that the expression of these two rubber transferases are regulated differently.[1]

Because it is unclear as to whether HRT1 contributes more to rubber production than HRT2 (or vice versa), we decided to include both enzymes in our construct. What we decided to *not* to include in our fusion protein but contained in the Latex Operon is the cDNA encoding small rubber particle protein SRPP, a coenzyme that is shown to have a positive role in facilitating rubber chain elongation via prenyltransferase activity.[4] The details of how exactly SRPP interacts with prenyltransferases at the molecular level is not well defined. In order to ensure that both the HRT1 and HRT2 proteins have access to SRPP, we do not want SRPP to be covalently linked to one or the other as part of the fusion protein. Instead, we will express SRPP via the Latex Operon, which expresses the rubber transferases and SRPP as separate proteins (the cDNA encoding for the corresponding proteins are separated by ribosome-binding sites). By doing so, the SRPP will be freely available for both the HRT1 and HRT2 components of *cis*-prenyltransferase.

In our final gene construct, an IPTG-inducible constitutive promoter (BBa_I712074) was placed at the beginning of the gene construct in order to regulate expression of the fusion protein. The Lpp-ompA gene sequence, which includes the (Gly-Ser-Ser-Ser)_2_ linker, was codon optimized for expression in *E. coli* K12 and synthesized as one single fragment. Because we were unable to acquire the assembled Latex Operon, the whole gene construct had to be reassembled from stock samples of synthesized gene fragments and inserted into a pSB1C3 backbone via Gibson assembly. In order to incorporate the two rubber transferases into our final construct, we proceeded to PCR amplify the sequences corresponding to HRT1 and HRT2 from the re-assembled Latex Operon. The HRT1 and HRT2 proteins are linked via a (Gly-Gly-Ser-Gly)_2_ linker. We inserted a double terminator (BBa_B0010 and BBa_B0012) at the very end of gene construct to ensure that the RNA polymerase does not leak through. In the end, a fusion protein

The substrates for the chain elongation reactions catalyzed by *cis*- prenyltransferase are isopentenyl-pyrophosphate (IPP) and dimethylallyl-pyrophosphate (DMAPP). These two isoprene monomers are already naturally produced in *E. coli* from the glycolytic products pyruvate and glyceraldehyde 3-phosphate (G3P) via two endogenous pathways – the mevalonate pathway and the non-mevalonate (MEP/DOXP) pathway.[5]

In order to upregulate the production of IPP and DMAPP, we decided to increase expression of the enzyme DXP synthase (DXS), which catalyzes the rate-limiting step of the MEP/DOXP pathway. The overexpression of DXS was achieved through transformation of the *E. coli* with the DXS gene construct (BBa_K2027040) developed by our predecessor team from 2016. As with the Latex Operon, we had to reassemble the DXS gene construct from synthesized gene stock samples and inserted the construct into a pSB2K3 plasmid backbone.

The next question we had to answer was whether IPP and DMAPP were capable of being transported out of the cell into the extracellular environment, since our polymerization machinery will be located on the cell surface. Although the membrane transport of terpenoids – the group of chemical compounds to which IPP and DMAPP belong – is not yet well characterized, previous research has suggested that ABC transporters facilitate the transport of endogenous secondary metabolites in plants across the membrane.[8] A recent study on phosphoantigen release in the immune system demonstrated that a specific ABC transporter regulates the extracellular release of IPP from dendritic cells.[2] Furthermore, based on data provided by a collaborator at Amyris, IPP will be observed in the extracellular matrix of *E. coli* when produced at high levels, indicating that there is some mechanism by which the isoprene monomers may be released into the extracellular environment. Although we were unable to find other information specifically regarding membrane transport of isoprenoids in *E. coli*, we hypothesized that IPP and DMAPP would be able to diffuse or be transported out of *E. coli* when both are overproduced via the MEP/DOXP pathway.

### Self-healing Materials Intro

Space presents challenging, harsh conditions for all materials. The constant threats of mechanical failure or fracture present challenging obstacles for long-term, sustainable space travel. When faced with the low-pressure vacuums and extreme temperature ranges experienced during missions, traditional glues lack the low-outgassing or low-volatility characteristics required for effective application in space (Thirsk et al., 2009). Traditional adhesives also lack intrinsic detection mechanisms that can uncover initial signs of material weakness, which usually begin with microscopic damage.

We designed a system of space-effective self-healing plastics that can address the need for material repair in space by applying principles from synthetic biology. *Bacillus subtilus,* a common soil bacterium with the longest current survival time in space (six years in endospore form aboard the NASA Long Duration Exposure Facility) has a natural capacity to produce the “super-glue” levan in the presence of sucrose (Horneck et al., 1994). Levan is a polysaccharide that contains no volatile organic compounds or other toxic emissions. Further, it is biodegradable and water-based, with no need for additional solvents. We thus used *B. subtilus*’ levan-producing pathway to create self-healing materials by embedding *B. subtilus* spores within a bulk plastic material. By inducing this pathway through light or oxygen activation, we can accurately control levan glue production. Specifically, if the material breaks or cracks, embedded *B. subtilus* would be exposed to oxygen and/or light and produce Levan’s glue to heal the material. The embedded bacteria will also sporulate under the anaerobic conditions of encapsulation, thus minimizing the need for sustained nutrient provision.

Due to the limited nutrient supplies available in space and the need to maximize the “shelf-life” of materials used in space missions, it was imperative to minimize the metabolic demand of *B. subtilis* used in our self-healing plastic. Thus, we began investigating viable routes for endospore encapsulation. As endospores, *B. subtilis* reduce their metabolic demand to negligible amounts by interrupting metabolic activities. This mechanism is triggered by unfavorable environmental conditions and is meant to promote survival of the bacteria. In fact, the bacteria can remain in this state for hundreds or thousands of years. Yet, upon studying the vegetative and sporulation cycles of *B. subtilis,* it became clear that very specific conditions were necessary to induce activation and germination. Specifically, these conditions included the requirement for nutrients germinants to be readily available. Encapsulation of free bacterial spores within the bulk of the material at issue was considered but rather quickly discarded. Such a method would require embedding a substantial amount of nutrients throughout the bulk material which could significantly impair its mechanical integrity.

This prompted the necessity to design an encapsulation system that would allow for evenly distributed nutrients access to the endospores at high concentrations and in specific locations. Thus, we started exploring options for microencapsulation. We sought an encapsulating material that fulfilled the need for feasible microencapsulation, was not toxic to bacteria, could be successfully embedded in a plastic bulk material and would break upon rupture of the bulk material. The method that most closely aligned with our aforementioned needs was agarose microencapsulation. Through an adaptation of a protocol described by Mathiowitz et al. 1999, we achieved viable encapsulation of *B. subtilis* within agarose microbeads. Agarose is a nonionic polysaccharide that naturally aggregates in water to form a gel and is thus an ideal polymer for encapsulation.

An essential component for a self-healing material is a method to detect breakage and in turn trigger the controlled secretion of an adhesive agent. We initially began exploring light activation as a means of controlled production of adhesive. We were primarily interested in the pDawn/pDusk system (outlined by Ohlendorf et al., 2012). pDawn and pDusk are a pair of plasmids that provide transcriptional regulation in response to light stimulus. pDawn induces the gene of interest in blue light-conditions, while pDusk induced the gene of interest in the absence of blue light. The first section our consolidated gene construct (BBa_K2485010) is analogous to the reversal step needed in pDawn, the optogenetic induction construct, to switch it from being deactivated in light conditions, to be activated in light conditions. We wanted to emulate the logic of pDawn in our design of an oxygen-sensitive construct. Fundamentally this meant having a promoter activated in anaerobic conditions that controls the transcription of a repressor, and having the associated promoter to that repressor controlling the transcription of the gene of interest, SPP. The first promoter, fnr, is active in anaerobic conditions, and produces a repressor for the second promoter, abrB. In aerobic conditions, as in when the material is broken, the repressor, *Spo0A*, will not be produced, and thus SPP, which results in levan production, will be generated.

Our constructs thus included BBa_K2485009, the *B. subtilis* fnr global anaerobic promoter with downstream *Spo0A* sporulation/repressor protein; BBa_K2485004, *B. subtilis* abrB promoter with downstream GFP (codon-optimized for *B. subtilis*); BBa_K2485006, the *Arabidopsis thaliana*-derived SPP enzyme for converting Suc-6-P to sucrose. Our consolidated construct BBa_K2485010 combines these individual constructs to create a controlled pathway for aerobic transcription of our gene of interest, SPP. Specifically, under anaerobic conditions, fnr promoter activation controls the transcription of the *Spo0A* protein repressor (and induces sporulation), which in turn inhibits the transcription of our gene of interest SPP via direct repression of the abrB promoter. Upon oxygen exposure, abrB inhibition is released, resulting in SPP expression, sucrose availability, and levan production.

If the material is intact, the anaerobically-induced promoted will result in generation of *Spo0A*, resulting in sporulation and no levan production. When the material is broken, the Bacillus no longer have induced production of *Spo0A*, and having a small amount of nitrate energy source surrounding them are now able to become vegetative cells. These cells then constitutively express SPP, and convert the sucrose-6’-phosphate in their substrate to sucrose, which they then convert to levan and energy for mass reproduction. When enough levan has been released to seal the crack, conditions again become anaerobic, and *Spo0A* is again produced, resulting in sporulation of the activated cells. This results in a minimal consumption of nutrients, and ability to break at a single site multiple times, as long as there remain sufficient nutrients. This is a substantial benefit over existing self-healing systems, which do not allow for repeated fracture at a site.

### BioBactery Intro

Currently, the ISS relies primarily on solar arrays for power, turning to lithium ion batteries when not in direct sunlight. But given the high costs of sending massive objects into space and the hazardous volatility of current batteries, these may not be the most sustainable sources for power. Looking for solutions in nature, we were inspired by the electric eel, whose linearly oriented electrogenic cells unipolarly pump ions in a single direction to generate an electric potential. This inspired our idea of a biobactery. Using *E. coli,* we mimicked the two key characteristics of eel cells: (1) stimulus-mediated unidirectional ion transport to generate a voltage and (2) linearly oriented cells that propagate this potential, like batteries in series. Ultimately, we aimed to use high-density *E. coli* colonies within a microfluidic device to polarly pump hydrogen ions via optogenetic activation of an ion channel, proteorhodopsin, to produce a voltage difference across the device.

We initially decided to use the naturally occurring polar auxin transport in plants to generate an electrical voltage to mimic the natural system perfected by electric eels. Plants have a naturally organized structure of aligned cells that display ion channels on only one side of the cells. This structure enables the unidirectional transport of auxins, plant growth hormones. We thought of exploiting this conveniently organized arrangement such that, upon stimulation, a concerted, unidirectional flow of ions would generate an electric potential. However, we quickly realized that we would not be able to express the engineered proteins in plants over the summer given the time for plants to germinate and the summer project timeline.

Following a thorough literature search, we found that previous research had shown evidence that bacterial cells have evolved to self organize into superstructures to unidirectionally pump ions and facilitate removal of toxic waste in highly-dense populations (HoJung et al. 2007). To further support this point, the researchers found that within these small chambers (width ~30 μm), the orientation of *E. coli*, their growth, and their motion is correlated to the chamber walls and location of chamber exits (HoJung et al. 2007). We saw potential in utilizing the self-organization, compact nature of high density *E. coli* colonies within a microfluidic chamber, and their rod-like morphology as the foundation for a new biological battery. Importantly, the suggested ability of *E. coli* to unidirectionally transport waste molecules was perfect in the context of our biological battery.

With the identification of a potential model system by which to create our battery, there were **three** main questions that had to be addressed. First and foremost, was it possible to engineer wild-type *E. coli* with a unipolar protein to generate a unidirectional flow of ions? Secondly, could we organize individual *E. coli t*o form a superstructure conducive to unidirectional ion flow? Finally, how much electric potential could the biobactery theoretically generate?

To create a voltage differential across the length of single *E. coli*, it was essential to localize our desired ion channel to a single pole within our organism. To accomplish this goal, it was first necessary to identify a single-cell organism capable of stable unipolar protein expression. Following an extensive literature review, we identified that the outer membrane protein IcsA of *Shigella flexneri*, was unipolarly expressed at the bacterial old pole (Goldberg et al. 1995, Robbins et al. 2001). This outer membrane protein is an autotransporter and is a member of the largest family of extracellular proteins in gram-negative bacteria (Pallen et al. 2003). It functions to mediate actin-based motility in *Shigella flexneri* and is essential for intra- and intercellular spreading through host epithelium (Goldberg et al. 1995, Bernardini 1989). Given the nature of *Shigella flexneri* mobility, IcsA localization is restricted to a single pole of the organism.

Having identified IcsA as potential candidate unipolar protein and having confirmed superstructured organization of *E. coli* in chambers, the next step was to investigate whether IcsA expression, localization, and stability could be maintained in *E. coli*. Further experimental investigations demonstrated that IcsA expression in *E. coli* was not only possible, but that the unipolar localization was maintained (Doyle et al. 2015). This finding, however, did not extend to *E. coli* strains that lack a complete lipopolysaccharides (LPS). This extracellular structure of most gram negative bacteria is composed of a saccharide region that is anchored to the outer membrane of bacteria by a lipid structure known as lipid A. Cells that have truncated LPS or lack the O antigen are no longer present with unipolar localization of IcsA (Sandlin et al. 1996, Van Den Bosch et al. 1997, Nikaido et al. 1987). Accordingly, it was essential to select a strain of *E. coli* that presented a complete LPS. We found that the *E. coli* strain W expressed a complete LPS while simultaneously satisfying the prescribed safety guidelines (Archer et al. 2011). With the validation that IcsA could be polarly expressed in our desired model organism, the next goal was to create a fusion protein between IcsA and an ion channel, to ensure that both were unipolarly localized to facilitate unidirectional ion flow. A deeper understanding of the structural features of IcsA, as well as its mechanisms of unipolar localization, was necessary to successfully engineer our fusion protein.

IcsA can be divided into three major domains. Beginning from the N-terminus: the signal sequence (amino acids (aa) 1-52), the passenger domain (aa 53-758), and finally a C-terminal translocation domain (aa 759-1102) (May et al. 2017). The N-terminal signal sequence mediates Sec-dependent transport of the protein across the inner membrane, the passenger domain functions to transport IcsA across the outer membrane, and the C-terminal translocation domain is necessary for export of IcsA (Brandon et al. 2003, May et al. 2017). Within IcsA, two different regions within the passenger domain were identified as sufficient for polar localization of the protein; the first region contained aa 1 to 104 and the second contained aa 507 to 620 (Charles et al. 2017). Of these two regions, it was found that aa 507-620, known as the cPT, was substantially more efficacious at polar location of IcsA. More importantly, polar protein localization using the cPT region of IcsA had been previously validated in *E. coli* (Doyle et al. 2015, Charles et al. 2017).

With the identification of the essential structural motifs required to localize IcsA to the old pole of *E. coli,* we began the process of engineering the fusion protein for unidirectional ion transport. To maximize the theoretical voltage differential of the biobactery, it was necessary to identify a protein that would allow for controlled ion movement. To further explain, if our ion channel of interest allowed for continuous, random ion movement there would not be unidirectional ion flow and no voltage differential could be produced. Accordingly, we began searching for ion channels with modular ion flow rates so that the ion channel could be inducibly opened or closed. To maximize the precision of ion channel activation, we sought to implement optogenetically controlled ion channels in our biobactery. Our first ion channels of interest were channelrhodopsins. However, it was found that functional expression of channelrhodopsins in *E. coli* is not possible due to the lack of glycosylation mechanisms within the model organism (Hou et al. 2012). We expanded our search into bacteriorhodopsins and proteorhodopsins with the hopes that these proteins could be expressed in *E. coli*.

While both of these rhodopsins could be synthesized within our model of interest, it is essential that bacteriorhodopsins form a hexagonal lattice structure in order to maintain their proton pumping functionality (Bratanov et al. 2015, Hou et al. 2011). We feared synthetic engineering of a bacteriorhodopsin to express the cPT region of IcsA would render the rhodopsin nonfunctional. Accordingly, we opted to use a proteorhodopsin, which is less dependent on the formation of its trimeric structure for ion movement (Yamashita et al. 2013). In fact, it has been shown that individual monomers contain all the necessary components to pump protons, albeit at a potentially lesser efficiency (Hussain et al. 2015). As a family, proteorhodopsins are a transmembrane proton pumping protein endogenous to SAR86 γ-proteobacteria. In response to light, proteorhodopsins pump protons out of the cell of its native organism to establish the proton motive force necessary to generate ATP (Bamann et al. 2014). Having identified the rhodopsin of choice, it is imperative that every *E. coli* transports its ions in same direction. While it is true that *E. coli* growing in high density colonies within chambers orient themselves in the same direction, if the old poles of the every *E. coli* are not facing in the same direction then, when activated, our engineered rhodopsin will mediate ion flow in opposite directions. If this occurs, the theoretical voltage differential produced will be reduced. Accordingly, after old pole localization of our engineered rhodopsins, it is essential that all old poles are facing in the same direction.

#### Voltage

The voltage protein we designed is a triple fusion protein between the proteorhodopsin, the sequence to localize unipolarly, and a fluorescent reporter. Its schematic is shown in Figure 1. Each domain has a distinct function. From the amino terminus to the carboxy terminus, these domains are the proteorhodopsin domain, the cPT domain, and the RFP domain. The first domain contains a green-light absorbing proteorhodopsin. We chose to use a proteorhodopsin rather than a channelrhodopsin or a bacteriorhodopsin, because eukaryotic channelrhodopsins are difficult to express in *E. coli* due to the requirement of post-translational modifications, and because bacteriorhodopsins form a hexagonal lattice structure which is very important to functionality, less so than the trimeric structure formed by proteorhodopsin (Hou et al. 2011, Yamashita et al. 2013).

**Figure 1:**
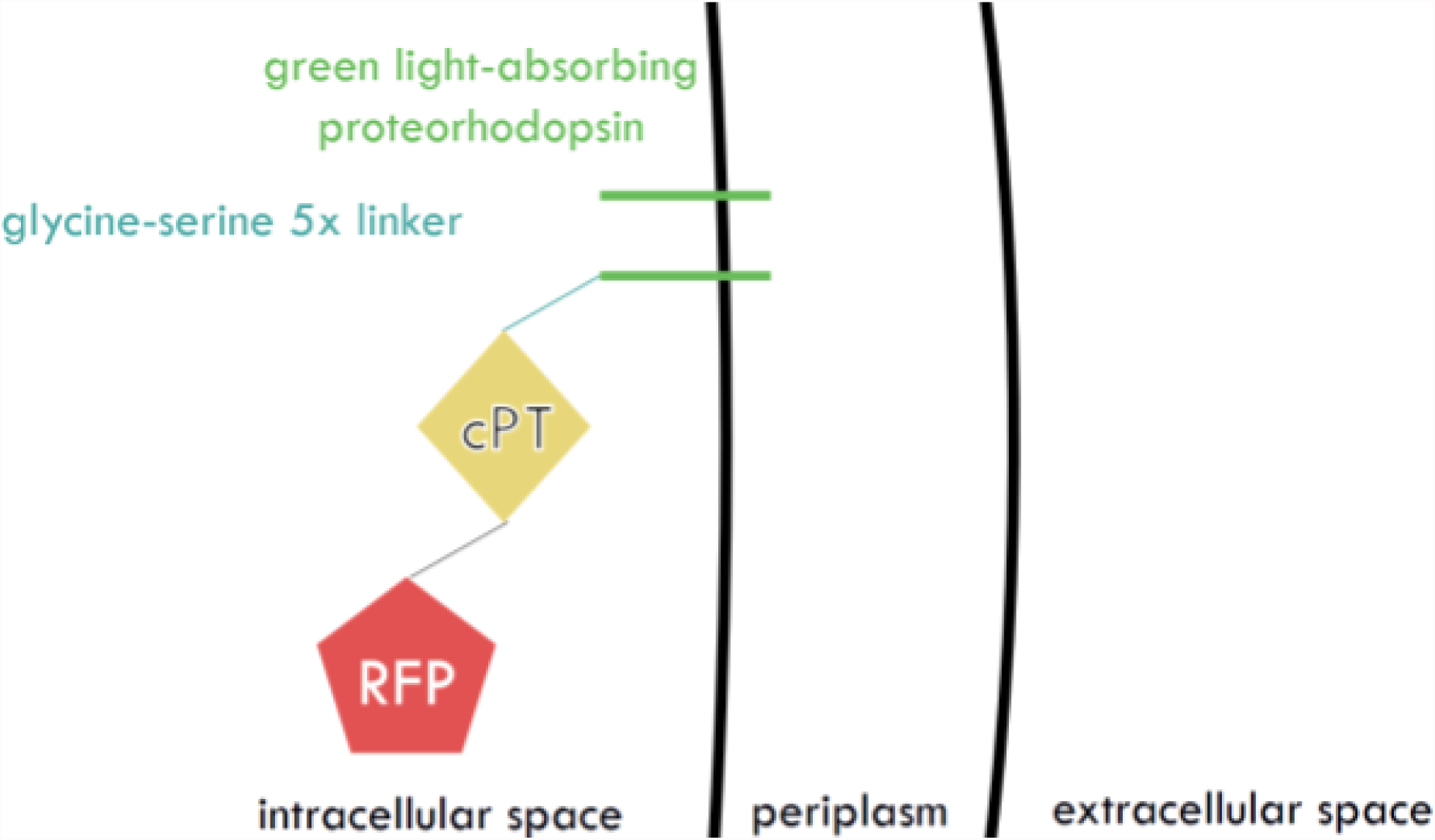
Schematic of the proteorhodopsin, cPT, RFP fusion protein. The proteorhodopsin is embedded in the inner membrane and linked to the cPT region with a GSx5 linker.

Proteorhodopsin is a transmembrane proton pumping protein endogenous to SAR86 γ-proteobacteria. In response to light, proteorhodopsin pumps protons out of the cell and helps its native organism establish the proton motive force necessary to make ATP (Bamann et al.). It does form a trimeric lattice structure in the inner membrane; however, it has been shown that the individual monomer contains all the necessary components to pump protons, albeit at a potentially lesser efficiency (Hussain et al. 2015). We sourced the sequence for the green-light absorbing proteorhodopsin from the 2012 Caltech iGEM team part BBa_K773002; however, codons towards the carboxy terminus were slightly modified to enhance GC content for IDT synthesis. This is the first domain in the protein because the carboxy terminus of proteorhodopsin is intracellular, while the amino terminus is extracellular, and the remaining domains must be intracellular for functionality (Stone et al. 2013).

The second domain of the voltage protein is the cPT, or central polar targeting, region of the larger IcsA protein. In its native organism, *Shigella flexneri*, the IcsA protein localizes to the outer membrane of the old pole and is responsible of the motility of the organism within mammalian host cells via actin polymerization. The cPT domain within this larger protein, when fused to GFP, has been shown to form a unipolar focus in 64% of *E. coli* cells (Doyle et al. 2015). It should be noted that this focus occurs intracellularly since neither the cPT domain, nor the GFP contain export signals.To fuse this domain to the carboxy terminus of the proteorhodopsin, we searched the literature for fusion proteins with proteorhodopsin. After being unable to find any previous attempts at this fusion, we decided to incorporate a flexible 5x Glycine-Serine linker between the proteorhodopsin and the cPT region.

The final domain of the voltage protein is a fluorescent reporter so that we can visualize the proteins localization. For this reporter, we chose to use FresnoRFP with an ex/em of 553/592 nm. We chose this protein because its ex/em is sufficiently far away from the excitation frequency of 525 nm of green-light absorbing proteorhodopsin, and we wanted to minimize excitation of the rhodopsin during visualization (Bamann et al. 2014). To link the cPT region to the RFP, we used the same sequence as was used by Dr. Marcia Goldberg, M.D. from Massachusetts General Hospital, who graciously gave us her plasmids which contained a cPT-GFP fusion. We showed that this fusion protein exhibited widespread unipolar localization in *E. coli*. In order to link cPT to GFP, the last four nucleotides of the cPT region were altered from “TCAT” to “ATCC.” We are unsure of the nature of this modification; however, it has been demonstrated to work efficiently.

By creating this fusion protein and demonstrating both the functionality of the proteorhodopsin and its unipolar localization, we hope to allow *E. coli* to generate a voltage differential across its length. Then, by aligning an entire culture of *E. coli* such that all the old poles, where the proteorhodopsin localizes, point in the same direction, each voltage will be additive like batteries in series. Although *E. coli* are very small, and proton diffusion is very fast, it has been shown that *E. coli* expressing proteorhodopsin will generate a voltage drop across the length of a cuvette in response to a unidirectional pulse of light (Sineshchekov et al. 2004). This is hypothesized to be a result of asymmetric excitation of the proteorhodopsin due to *E. coli* cells acting as a lens, focusing light on the far end of the cell. By controlling the expression of the proteorhodopsin to be unipolar, we hope to be able to generate this same sort of voltage without the need for the light to be unidirectional.

#### Orientation

We first considered and tested a chemotactic approach, where, in theory, the *E. coli* would swim toward a chemoattractant and in the process orient along their old pole. We tested whether LB served as a chemoattractant for *E. coli* in minimal media in our microfluidic device, but unfortunately observed random movement of our bacteria. Upon further literature research, we found that due to the intrinsic nature of *E. coli* locomotion, at any given time point the orientation of a single *E. coli* would vary greatly even if the net movement was towards the chemoattractant. In fact, many genera of motile bacteria, including *E. coli*, engage in a behavior called swarming when surrounded by a dense colony in a bulk fluid (Darnton et al. 2010). During swarming, *E. coli* continually reorient themselves by random jostling with their neighbors, randomizing their directions within a few tenths of a second: for example, the cells may move laterally if they collide with adjacent cells or may reverse their direction abruptly as the motors that actuate the flagella change direction (Akbarzadeh et al. 2012). Overall, the cells tend to separate from each other principally due to their difference in speeds or rotational diffusion. Thus, relying on the bacteria’s natural mode of locomotion in response to a stimulus were not feasible options for our desired outcome. More importantly, given limitations in the production of microfluidic devices, we quickly realized it would not possible to craft a channel that could maintain a concentration gradient over time. Without a concentration gradient, the theory supporting this hypothesis was flawed and we abandoned chemotaxis mediated orientation.

We then identified a mechanical solution in order to impose the desired highly-organized and super-oriented alignment. Specifically, we functionalized the cells with superparamagnetic microbeads and employed an applied, external magnetic field to induce the mechanical rotation of the cells towards the magnet, as determined by the magnetic force on the beads. Superparamagnetic microbeads were selected for this application due to their ability to increase in magnetization when a magnetic field is applied, and lose their magnetic properties upon removal of the field (Akbarzadeh et al. 2017). This implies that no residual magnetization is observed, eliminating the potential risk of the beads interfering with the cells’ biological functions and behavior. From a literature review, we were able to determine that previous studies had shown the successful integration of a biotinylation site within the coding sequence of IcsA without disrupting or interfering with its function (May et al. 2012). Through the integration of a biotinylation site on the IcsA aa 87, we were able to biotinylate the cells and subsequently enable the formation of biotin-streptavidin bonds with streptavinated paramagnetic microbeads in solution. The biotin-streptavidin bond, is one of the strongest found in nature, thus further confirming the stability and reliability found in the choice of this *induced orientation* method.

In order to properly orient our *E. coli*, we needed to find a unipolarly localizing protein that could be mechanically manipulated to orient the *E. coli*, which we found in IcsA. The IcsA protein additionally consists of two parts; the passenger domain, or the region responsible for unipolar localization, and the beta barrel, the region responsible for actin polymerization in the endogenous protein (May et al.). IcsA has also been shown to be able to host a biotinylation site within its coding sequence without changing the functionality of the protein. This further makes IcsA an ideal candidate for the orientation construct because if the protein is biotinylated and localizes the old pole on the outer membrane, cells could be washed with streptavidin coated magnetic beads, allowing for a biotin-streptavidin bond between the cell and the magnetic bead. The *E. coli* would then be able to be oriented by applying a magnetic field in liquid culture.

In our initial design of the orientation construct, we designed the coding sequence to be a conjugation between the IcsA passenger domain and GFP using 5X poly-glycine linker. A biotinylation site was encoded at amino acid 87, which has been shown to not disrupt the functionality of IcsA (May et al.). We used the IPTG inducible promoter from the Parts Registry, BBa_R0011, to control the expression of our construct. We used the ribosome binding site from the Parts Registry, BBa_B0034, to allow for translation of the protein. After the coding sequence of the protein, we included two terminators, BBa_B0010 and BBa_B0012.

However, upon further literature research, we found that when the IcsA passenger domain has been conjugated to GFP, the protein no longer localizes to the outer membrane of *E. coli*, but instead to the intracellular space on the old pole of the cell (Charles et al.). To amend this flaw in the construct design and ensure unipolar localization to the outer membrane of the old pole of *E. coli*, we then planned to take out the GFP sequence and clone the beta barrel sequence back into the plasmid, as the full sequence of IcsA has been shown to localize to the old pole of *E. coli*. The beta barrel sequence was then also ordered as a secondary addition to the main orientation construct.

We also designed a construct to enhance biotinylation of our orientation construct to ensure that the biotinylation site on the IcsA protein would be biotinylated, as it needs to be able to bind strongly to the streptavidin-coated magnetic beads. To accomplish optimal biotinylation of IcsA, we designed our BirA construct. BirA is the protein responsible for biotinylation endogenously in *E. coli*, and acts by associating with biotin to form the holobirA complex, which then associates with the apoBCCP complex to transfer biotin in conditions of high biotin demand (“Bifunctional ligase/Repressor BirA”). For our biotinylation construct, we began with the Biobrick Prefix, followed by a rhamnose inducible promoter from the Parts Registry, BBa_K914003. We next used the ribosome binding site from the Parts Registry, BBa_B0034, to allow for translation of birA. The coding sequence for birA, obtained from the NCBI database, was codon optimized for *E. coli* and attached after the ribosome binding site, along with a 6X poly-His tag to allow for protein visualization. We then added the Parts Registry standard double terminator, BBa_B0015, followed by the Biobrick suffix.

## Materials and Methods

The Materials and Methods section should provide enough detail to allow suitably skilled investigators to skilled investigators to fully replicate your study. Specific information and/or protocols for new methods should be included indetail. should be included in detail. If materials, methods, and protocols are well established, authors may cite aricles where those protocols where those protocols are described in detal, but the submission should include sufficient information to be understod to be understood independent of these references.

Protocol documents for clinical trials, observational studies, and other **non-laboratory** investigations may bemay be uploaded as suporting information. <u>Read the suporting information guidelines guidelines</u> for formatting instructions. We recommend depositing **laboratory protocols** at <u>protocols.io.</u> Read detailed <u>instructions for depositing and sharing your laboratory protocols</u>.

- Large data sets, including raw data, may be deposited in an apropriate public repository. <u>See our list of recommended repositories.</u>
- Methods sections describing research using cell lines must state the origin of the cell lines used cell lines used. <u>See the reporting guidelines for cell line research for more information</u>.

### Laboratory Protocols

To enhance the reproducibility of your results, we recommend and encourage you to deposit laboratory protocols in <u>protocols.io,</u> where protocols can be assigned their own persistent digital object identifiers (DOIs).

To include a link to a protocol in your article:

1. Describe your step-by-step protocol on protocols.io
2. Select **Get DOI** to issue your protocol a persistent digital object identifier (DOI)
3. Include the DOI link in the Methods section of your manuscript using the following format provided by protocols.io: http://dx.doi.org/10.17504/protocols.io.[PROTOCOLDOI]

### BioRubber Materials and Methods

Both NEBuilder® HiFi DNA Assembly Master Mix (Gibson assembly) and GeneArt® Seamless Cloning and Assembly Kit were both used for gene assembly. For cloning purposes, transformations were performed on NEB®5α Competent *E. coli*.

Once all the relevant gene cloning and assembly was completed, we proceeded to do a series of transformations on NEB® T7 Express Competent *E. coli* for the purpose of protein expression:

We attempted to do a triple transformation of plasmids containing the ompA+*cis*-PT construct, the DXS construct, and Latex Operon into T7 cells. Unfortunately, after multiple failures with triple transformation, we were only able to successfully perform a double transformation of the ompA+*cis*-PT construct (in pSB1C3) and DXS construct (in pSB2K3). Because these double transformants do not contain the Latex Operon, SRPP will not be expressed. Due to the absence of SRPP, we expected a down-regulation of *cis*-prenyltransferase activity and a consequent decrease in the amount of rubber produced by these double transformants. We also performed a double transformation of the Latex Operon (in pSB1C3) and DXS (in pSB2K3) into another group of T7 cells. This group was to be used for comparison purposes in the latex production assay.

These two groups of double-transformed T7 *E. coli* were cultured separately in LB media containing chloramphenicol and kanamycin. Once the bacterial growth in the liquid cultures reached logarithmic phase, reagents for induction of gene expression were introduced.

For induction of latex production, IPTG (500 mM), MgSO_4_ (500 mM), and glucose (1 M) were directly added to the liquid cultures, which had been growing at 37 °C. Protein expression of the DXS, ompA+*cis*-PT, and Latex Operon constructs are controlled by an IPTG-inducible promoter, so the introduction of IPTG will activate transcription. Magnesium ion (Mg^2+^) is an important cofactor for the catalytic activity of HRT1 and HRT2, as it influences the affinity of *cis*-prenyltransferase for its IPP substrate.[3] The addition of MgSO_4_ to our liquid cultures will increase the efficiency of HRT1 and HRT2 in using IPP substrate to initialize the polymerization of *cis*-1,4 polyisoprene. Glucose is provided to the T7 *E. coli* as the substrate for glycolysis, which produces the pyruvate and glyceraldehyde 3-phosphate (G3P) that are converted into IPP and DMAP via the MEP/DOXP pathway.

Five days after induction, an extraction protocol (courtesy of Gordon Sun, 2016 S-B iGEM) was performed on both cultures of double-transformed cells. Centrifugation of the cell samples yielded two fractions – a cell pellet and an LB-media supernatant. Chloroform is added to the suspended cell solution to lyse the cells and dissolve any polyisoprene produced. After filtering the suspended cell solution, the filtrate, which contains only chloroform and its solutes, was treated with methanol. The addition of methanol (polar) will precipitate any polyisoprene (non-polar) contained in the filtrate. A similar procedure was used to precipitate any polyisoprene in the LB-media supernatant.

The resultant precipitant, a flaky white polymer suspended in methanol, was dried overnight at 40 degrees C. To definitively identify the contents of the polymer material, the dried extracts were sent for chemical analysis, including molecular weight analysis, ^1^H-NMR, and ^13^C-NMR (courtesy of David Shintani, University of Nevada, Reno).

### Self-Healing Materials and Methods

To verify that *B. subtilis*-produced Levan could be used as a space-effective adhesive, we conducted a series of tensile tests to determine whether Levan could be applied towards a wide range of materials with a strength comparable to commonly-used glues. Our mechanical tests were conducted at Stanford’s Soft & Hybrid Materials Facility with the Instron 5565 tension tester. The adhesive strength of Levan produced from *B. subtilis* (following the protocol by Abdel-Fattah et al., 2005 & Abou-Taleb et al., 2014) was compared against Elmer’s glue (synthetic polyvinyl acetate glue), and silicon epoxy glue. Tested materials included cardboard, rubber, and actylonitrile butadiene styrene (ABS) plastic. Adhesive strengths were measured by tensile strength of a material until mechanical failure.

Our constructs were ordered from IDT and inserted into the empty pBS1c vector (integration at amyE locus) via restriction ligation. Between amyE loci, pBS1c contains flanking homology regions, a cat resistance cassette for chloramphenicol selection in *B. subtilis*, and a MCS containing a rfp-cassette flanked by the six BioBrick restriction sites (EcoRI, NotI, XbaI (upstream); SpeI, NotI, PstI (downstream)). Downstream of the backbone’s ori is a bla resistance gene allowing for ampicillin resistance selection in DH5α *E. coli* transformations. Cloning with BioBrick-standard restriction ligations into DH5α *E. coli* was screened via selection for white colonies, indicating successful removal of the rfp-cassettes, while unsuccessful cloning leads to formation of red colonies in *E. coli*. Plasmids were then miniprepped and transformed into super-competent *B. subtilis*.

To test the functionality of our SPP gene construct, we transformed *B. subtilis* with our BBA_K2485006 plasmid and used ammonium molybdate assays based on “Assay of Inorganic Phosphate, Total Phosphate and Phosphatases” (Ames, 1956) to test rates of sucrose-6-phosphate (S6P, provided in phosphate solution) to sucrose conversion in SPP-transformed *B. subtilis*. When converted, S6P releases inorganic phosphate ions, which react with ammonium molybdate in solution to produce a deep-blue colored complex. These assays allow us to determine SPP enzyme functionality when transformed into *B. subtilis* by quantifying rates of sucrose production from provided S6P. This conversion induces controlled levan production, as *B. subtilis* have an endogenous capacity to produce levan polymer in the presence of sucrose and not sucrose-6-phosphate.

In our first iteration, we used a 10 mL volume 1A976 super-competent (s.c.) *B. subtilis* cultures with a short S6P incubation period (5 minutes at room temperature) on lysed *B. subtilis* cells for our assay. Lysed 1A976 s.c. *B. subtilis* cultures (not transformed) served as our control condition; lysed SPP-transformed 1A976 s.c. *B. subtilis* cultures as experimental. OD-820 measurements were taken of final phosphate and ammonium molybdate-incubated solutions, with our control measurements serving as the blank baseline measurement. With our second iteration, we used 25 mL volumes of control 1A976 s.c. *B. subtilis* cultures and experimental SPP-transformed 1A976 s.c. *B. subtilis* cultures. Here, we tested both whole-cell and lysed-cell conditions for both control and experimental cultures. Thus, the assay was performed with four total *B. subtilis* samples: 1) control 1A976 s.c. *B. subtilis,* intact; 2) control, lysed; 3) SPP-transformed 1A976 s.c. *B. subtilis*, intact; 4) SPP-transformed, lysed, to compare rates of enzymatic activity between whole-cell and lysed *Bacillus.* Further, we increased our S6P incubation time to 1 hour at 37°C. Remaining protocol steps (namely solution preparation) were consistent with our first iteration.

With in-depth understanding of *B. subtilis*’ metabolic demands and vegetative-endospore cycle, we concluded that it was necessary to encapsulate the bacterial spores in nutrient-filled microbeads. Having evaluated diverse routes of encapsulation, we discarded those that only allowed for one-time rupture. For example, core microencapsulation refers to encapsulation mechanisms that generate a hard shell surrounding a liquid core. Such a method would have provided an aqueous environment which, in the presence of nutrients, approximates the requirements for spore germination. Additionally, a liquid core was problematic in the fact that it constituted a design limitation: if the microbeads are composed of liquid cores, their contents are lost upon the first rupture. Such a design did not meet our objective of creating a renewable, sustainable, self-healing material. Thus, we opted for methods of matrix encapsulation, where the encapsulating material and the encapsulated agent are mixed in a homogenous matrix, which can thus be ruptured repeatedly.

The material of choice for our encapsulation was low melting point agarose. By exploiting the large temperature difference between the gelation and melting temperature of agarose, we noticed that agarose microbeads could form gels at low temperatures (~15°C) and once formed do not melt until 70°C. This enabled us to synthesize sucrose-loaded agarose microbeads with encapsulated endospores, which could feasibly be incorporated in the molten bulk material (PCL), since its melting point is below agarose’s melting point. The general steps of our agarose microbead encapsulation protocol is as follows:

1. Completely dissolve low-gelling-temperature agarose in water at elevated temperatures,
2. Cool it to physiological temperatures (~37°C) without the microbeads forming a gel
3. Add the endospores and the sucrose to the solution
4. Add mineral oil to yield a water-in-oil emulsion
5. Drastically cool the solution to below agarose gelation point (~5°C) to form gel microbeads. Our “Agarose Microbead Encapsulation” protocol is adapted from Hempel, K, et al.’s “A simple mechanical procedure to produce encapsulated cells” to encapsulating *Bacillus subtilis* spores into agarose microbeads (Gu 2017).

### BioBactery Materials and Methods

All PCR reactions for cloning were done using NEB Q5 or NEB OneTaq, with the protocol provided on the NEB website with annealing temperatures determined by the NEB Tm calculator (“Q5® High-Fidelity DNA Polymerase,” “OneTaq® DNA Polymerase”). All gels were run at 100V for 50 minutes using a 1% agarose gel. To gel extract or PCR cleanup, we used the Zymogen DNA Clean & Concentrator kit and the Zymoclean Gel DNA Recovery Kit (“DNA Clean & Concentrator™-5,” “Zymoclean™ Gel DNA Recovery Kit”). For DNA assembly we used NEB EcoRI - HF and NEB PstI - HF for digestion and T4 DNA ligase for ligation, and the NEB Gibson Assembly Master Mix or GeneArt Seamless Cloning and Assembly (“Gibson Assembly® Master Mix,” “GeneArt® Seamless Cloning & Assembly”). To isolate plasmid DNA, we used the Zyppy Plasmid Miniprep Kit (“Zyppy™ Plasmid Miniprep Kit”). We used Elim Biopharmaceuticals for our sequencing needs (“ELIM BIOPHARM”).

We used AlexaFluor 405 streptavidin conjugate for the visualization of our biotinylated IcsA construct (“Streptavidin, Alexa Fluor 405 conjugate”), a generous gift from Thermo Fisher Scientific. We used streptavidin coated magnetic beads from BioLegend to validate the magnetic activity of the *E. coli* (“MojoSort™ Streptavidin Nanobeads”). We used the Zeiss AxioImager Z1 light microscope and the Evos XL Digital Inverted Microscope for microscopy (“Axio Imager 1 for Life Science Research,” “EVOS® Digital Microscopes”).

The microchannel devices were designed in SolidWorks. The design (see Figure 1) consisted of two wells (inlet well and outlet well) of diameter 2mm for the placement of probes to measure electric potential and for the insertion of *E. coli*. The two wells were connected by a microchannel (width 150 um). The devices were prototyped with a biocompatible silicone elastomer, PDMS (Sylgard 184, Dow Corning, Midland MI). The channel structure of the chips was formed by soft lithography: a negative master mold for the the channels was fabricated with a UV-curable epoxy (SU8 by MicroChem, Newton MA) by conventional contact lithography. Liquid PDMS pre-polymer, in a 10:1 ratio of catalyst and resin, was vigorously mixed, degassed in a vacuum chamber and poured onto the mold to a thickness of about 2 - 3 mm and cured in an 80*°C* oven for 90 minutes. Following the curing step, the elastomer was peeled off and cut into individual chips. The holes at the end of each channel well punched using a 2 mm luer stub. The chips were then baked for 30 more minutes at 150*°C* to increase hardness and Young’s modulus of the PDMS, to ensure that the micrometer-scale features were more stable against spontaneous collapse. The chips were then sealed to a #4 microscope cover glass by plasma wand high radio-frequency bonding. Before use, the channels were primed to increase hydrophilicity and reduce surface tension. The priming involved overnight incubation in a solution of 17:2:1 nuclease-free diH_2_O: Tween-20 (0.1%): BSA.

## Results, Discussion, Conclusions

These sections may all be separate, or may be combined to create a mixed Results/Discussion section (commonly labeled “Results and Discussion”) or a mixed Discussion/Conclusions section (commonly labeled “Discussion”). These sections may be further divided into subsections, each with a concise subheading, as appropriate. These sections have no word limit, but the language should be clear and concise.

Together, these sections should describe the results of the experiments, the interpretation of these results, and the conclusions that can be drawn.

Authors should explain how the results relate to the hypothesis presented as the basis of the study and provide a succinct explanation of the implications of the findings, particularly in relation to previous related studies and potential future directions for research.

PLOS ONE editorial decisions do not rely no perceived significance or impact, so authors should avoid overstating their conclusions. See the <u>PLOS ONE Criteria for Publication</u> for more information.

### BioRubber Results/Discussion

Approximately 0.014 g of dried polymer material was extracted from the cell solution of double transformants containing the plasmids with latex operon and ompA. Results from molecular weight analysis showed that the material in this sample had a weight-average molecular weight of Mw = 2.2468 × 10^5^ g/mol (see appendix files 1 and 2). This is an acceptable molecular weight value for cis-1,4 polyisoprene, although high quality natural rubber typically has a molecular weight of Mw > 1.0 × 10^6^ g/mol. However, when H-NMR was performed on the sample, the peaks observed in our sample spectrum did not match the peaks in the standard spectrum for cis-1,4 polyisoprene (see appendix file 3). Further research needs to be done expressing this material in larger quantities to test its physical properties.

In order to kickstart the process of strain optimization, we modeled our biosynthesis pathway using elementary flux mode analysis (EMA). Elementary flux modes (EFMs) are a concept used to analyse metabolic networks. An EFM is defined formally as the “characteristic (support-minimal) vectors of the flux cone that contains all feasible steady-state flux vectors of a given metabolic network” (Klamt 2017). In other words, an EFM is essentially a representation of possible ways to use carbon, nitrogen, and other inputs to make metabolic products used to keep the organism “alive”, or our desired product, cis-1,4 polyisoprene. Additionally, by allowing us to visualize what our theoretical yields might be, EFM analysis highlights possible knockouts of nonessential metabolic pathways that compete with our rubber biosynthesis pathway.

An analogy may be useful to describe our logic. Suppose you have 10 bags of flour, 5 pounds of butter, and 10 bags of sugar, and predefined measurements of a few other miscellaneous ingredients (e.g. chocolate chips, baking soda, sprinkles, icing). What are all the possible baked goods you can make with that set of ingredients? Are there certain combinations that are not as useful, and can we knock those recipes out of your recipe book so you no longer make them?

All modeling is constraint-based, and is calculated solely on stoichiometric relationships. If given more time, we would also like to explore network embedded thermodynamic (NET) analysis to model intracellular versus extracellular polymerization, and vary inputs into system (e.g. pH, temp, kinetics).

First, we wanted to explore if the mevalonate or non mevalonate pathway had higher theoretical yields. Both pathways can be used to make rubber, but which has higher yields? The non-mevalonate pathway displayed higher theoretical yields. Thus, we chose to employ this endogenous pathway for our rubber-producing strain.

**Figure 3.**
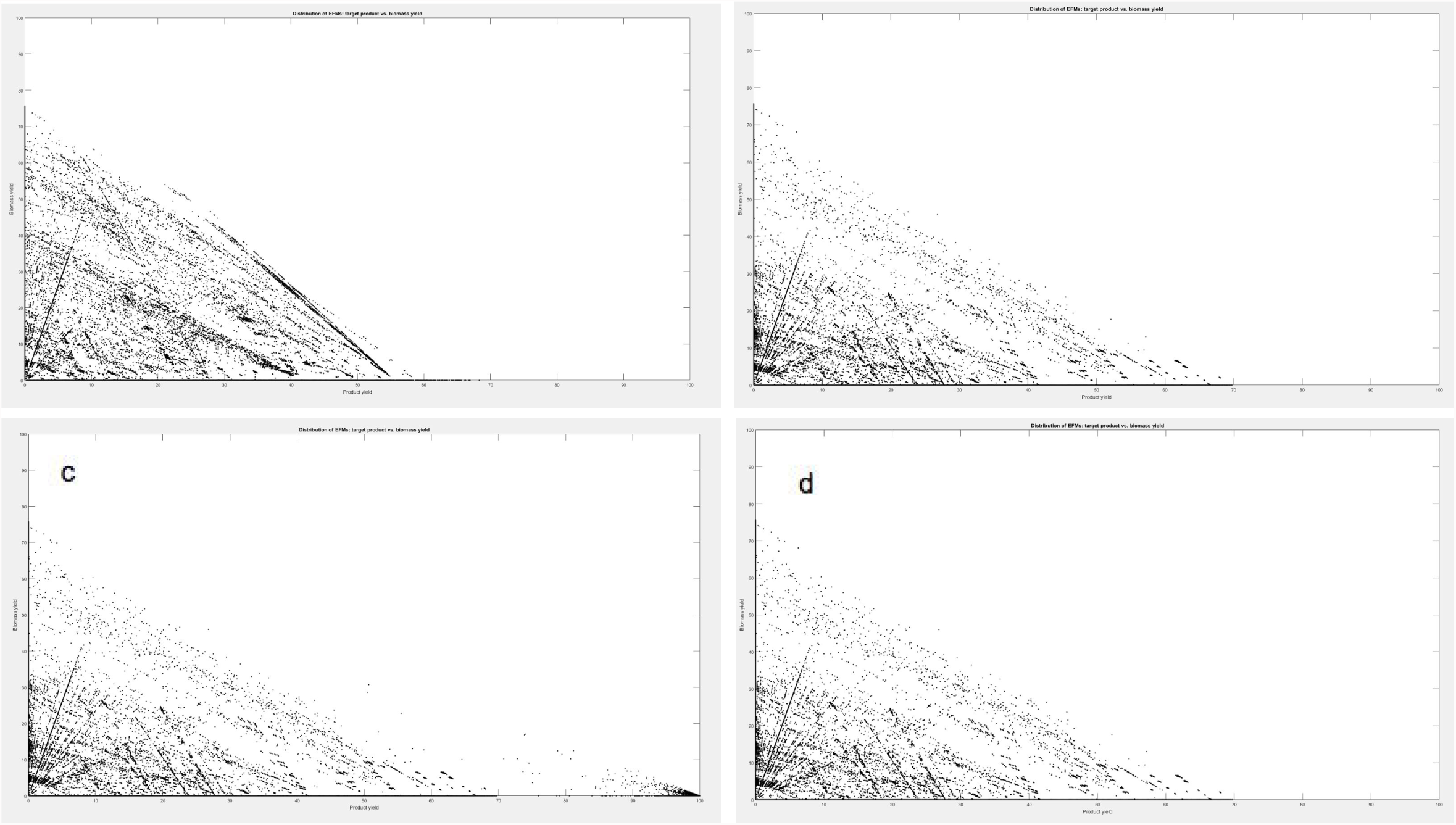
Each dot represents an EFM. X axis: Product yield as % [C-mol]; Y axis: Biomass yield as % [C-mol]. **a.** *Non-Mevalonate Pathway:* Rubber max with Biomass Production: 68.1% [C-mol/C-mol]; Rubber max without Biomass Production: 69.8% [C-mol/C-mol] **b.** *Mevalonate Pathway:* Rubber max with Biomass Production: 58.1% [C-mol/C-mol]<br/>Rubber max without Biomass Production: 68.3% [C-mol/C-mol] **c.** *Non-Mevalonate Pathway, Isoprene Feed:* Rubber max with Biomass Production: 99% [C-mol/C-mol] Rubber max without Biomass Production: 100% [C-mol/C-mol] **d.** *Non-Mevalonate Pathway, Glucose + Glycerol Feed:* Rubber max with Biomass Production: 68.1% [C-mol/C-mol] Rubber max without Biomass Production: 69.8% [C-mol/C-mol]

Second, we learned that our proposed recycling method appears promising by modeling glucose and glycerol vs. isoprene feed. If given an isoprene feed from recycled rubber (instead of glucose + glycerol carbon source), our *E. coli* could theoretically have 100% yield of rubber product.

### BioRubber Appendix files

1. http://2017.igem.org/wiki/images/4/42/T--Stanford-Brown--mwrub.pdf
2. http://2017.igem.org/wiki/images/6/69/T--Stanford-Brown--mwrub2.pdf
3. http://2017.igem.org/wiki/images/7/76/T--Stanford-Brown--stdrubmw.pdf

### Self-Healing Materials Results

We conducted a series of tensile testing experiments to determine the adhesive strength of B. subtilis-produced Levan 1) in comparison to traditional glues and 2) in determining potential applications to a variety of materials. Mechanical tests were conducted at Stanford’s Soft & Hybrid Materials Facility with the Instron 5565 tension tester. We measured an adhesive strength of 2.082 MPa for Levan; 0.842 MPa for Elmer’s glue; and 14.626 MPa for silicon epoxy glue (Fig. 1). Levan thus compares well and can serve as an effective alternative to traditional consumer glues (about 2.47x stronger than Elmer’s glue). Further, we found that Levan works best on cardboard (3.342 MPa) and plastic (2.002 MPa) material surfaces, and has a lower adhesive strength when applied to rubber (0.952 MPa) (Fig. 2).

**Fig. 1:**
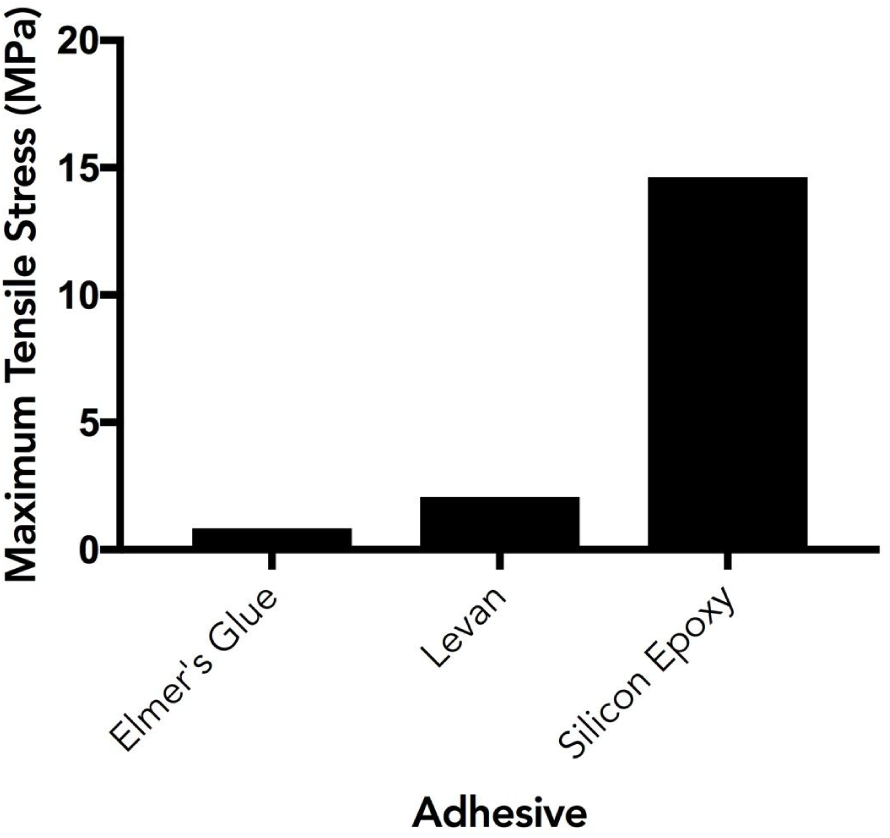
Axial Loading Capacity of Varying Adhesives on ABS.

**Fig. 2:**
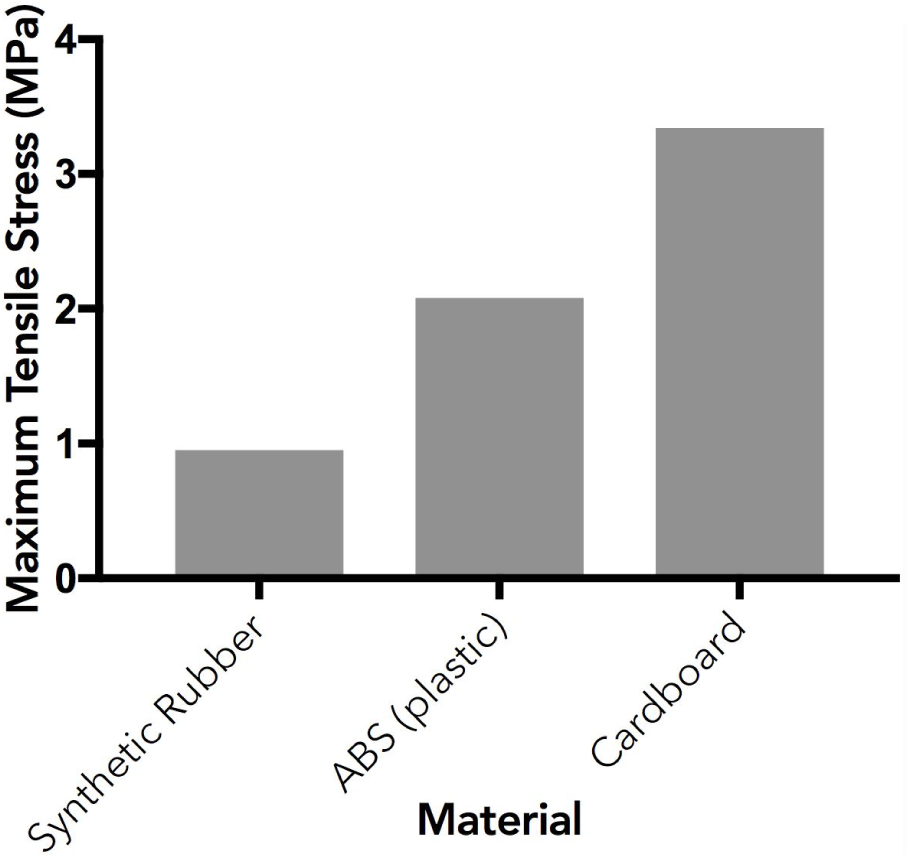
Axial Loading Capacity of Levan on Varying Materials.

For our construct testing, we inserted our gene of interest, SPP (allows for controlled production of Levan by converting provided “inactive” sucrose-6-phosphate to “active” sucrose, thus triggering *B. subtilis* Levan synthesis), into the empty pBS1c backbone with BioBrick-standard restriction ligations into DH5α *E. coli*. Plasmids were then miniprepped and transformed into 1A976 super-competent *B. subtilis*. Colony PCR and sequencing results showed successful insertion for SPP (amplification on all 20 picked colonies with SPP-specific primers, gel pictured below).

**Fig. 3:**
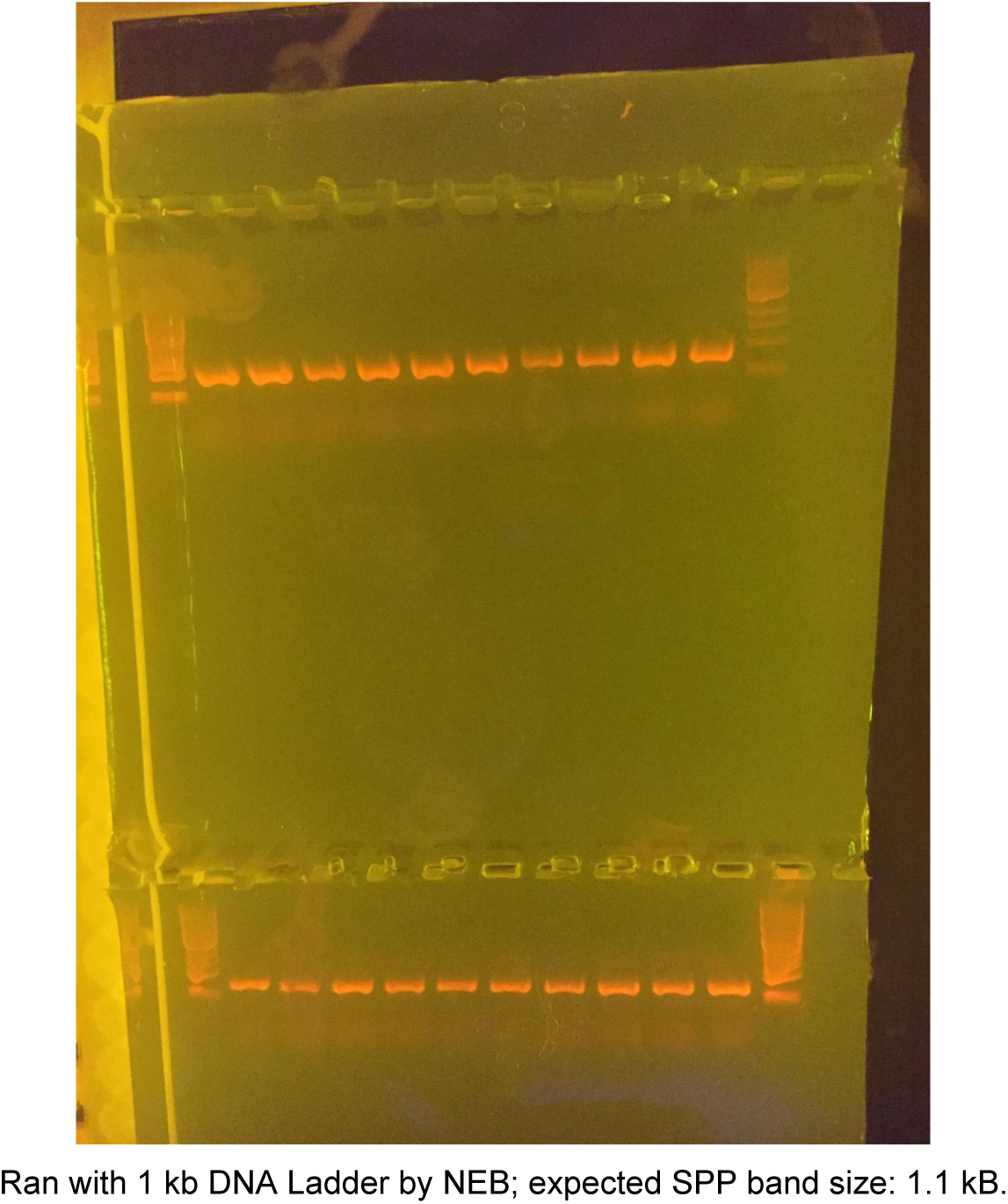
SPP Transformation with NEB’s 1 kb DNA Ladder

Following successful SPP transformation into *B. subtilis*, we conducted a series of ammonium molybdate assays to test rates of sucrose-6-phosphate (S6P, provided in phosphate solution) to sucrose conversion in SPP-transformed *B. subtilis*. These assays allow us to determine SPP enzyme functionality when transformed into B. subtilis by measuring rates of sucrose production from external S6P. This conversion confers our system’s levan-producing capacity, as *B. subtilis* have an endogenous capacity to produce levan polymer simply in the presence of sucrose. From our first ammonium molybdate assay iteration (using 10 mL *B. subtilis* culture volumes; lysed cell conditions; 5 minute at RT S6P incubation), we measured an OD820 reading of 0.055 on our SPP-transformed *B. subtilis*, blanked on the control *B. subtilis* (Fig. 4). This corresponds to a conversion of 0.56% of S6P in solution despite the low culture volumes, insufficient incubation period, and lower temperature conditions (10 mL cultures; 5 min at RT incubation periods against suggested 20 min at 45° or 1 hour at 37°C).

**Fig. 4:**
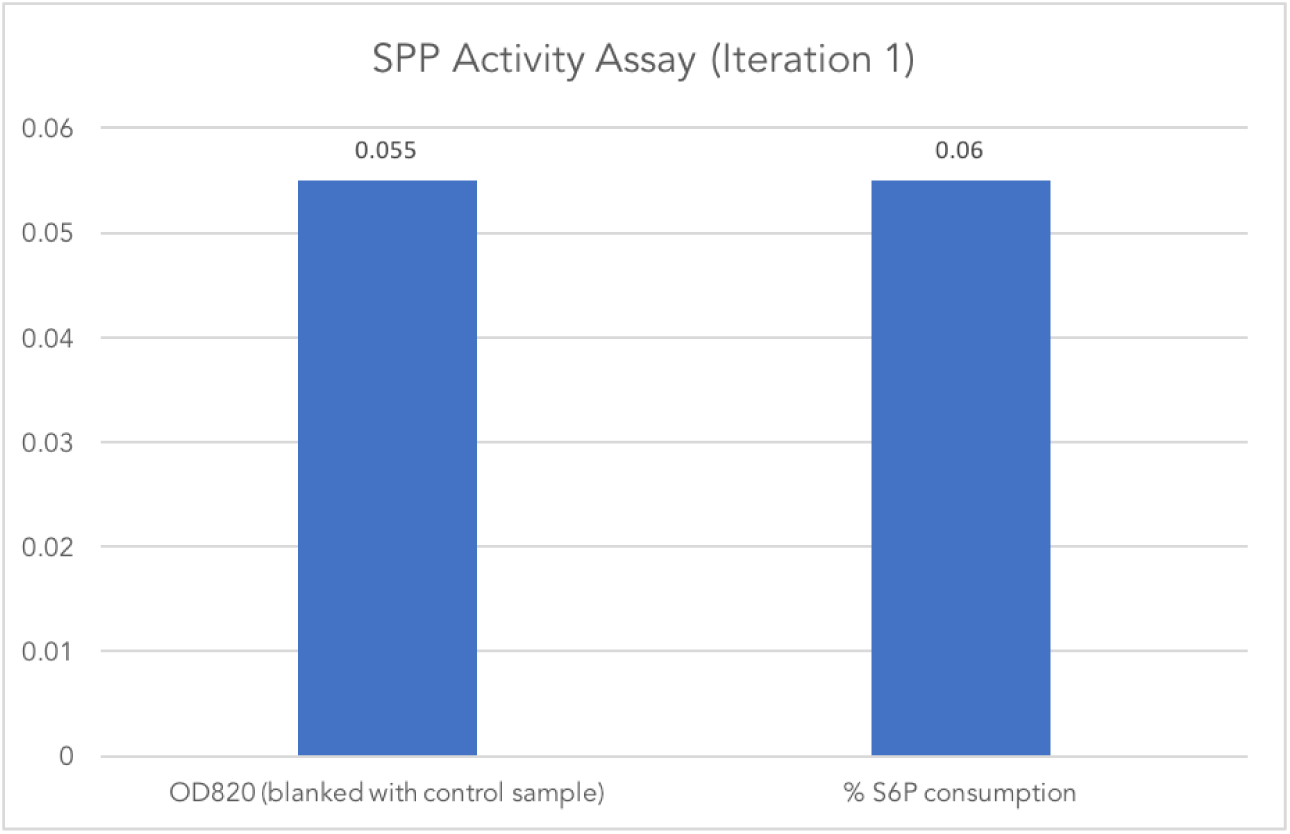
Ammonium Molybdate Assay with 0.56% (lysed-cell) S6P to Sucrose Conversion in SPP-transformed 1A976 *B. subtilis*

For our second iteration (using 25 mL *B. subtilis* culture volumes; both whole-cell and lysed cell conditions; 1 hour at 37°C S6P incubation), we measured an OD820 reading of 0.790 on intact-cell conditions and 1.157 on lysed conditions, both blanked on control *B. subtilis* (Fig. 5). This corresponds to a conversion of 8.04% and 11.8% of S6P in the intact-cell and lysed Bacillus, respectively. Intact cells perform comparably to (but predictably not as well as) lysed cells for S6P to sucrose conversion. These results were obtained with larger culture volumes, longer incubation periods, and higher incubation temperatures (25 mL cultures; 1 hour at 37°C).

**Fig. 5:**
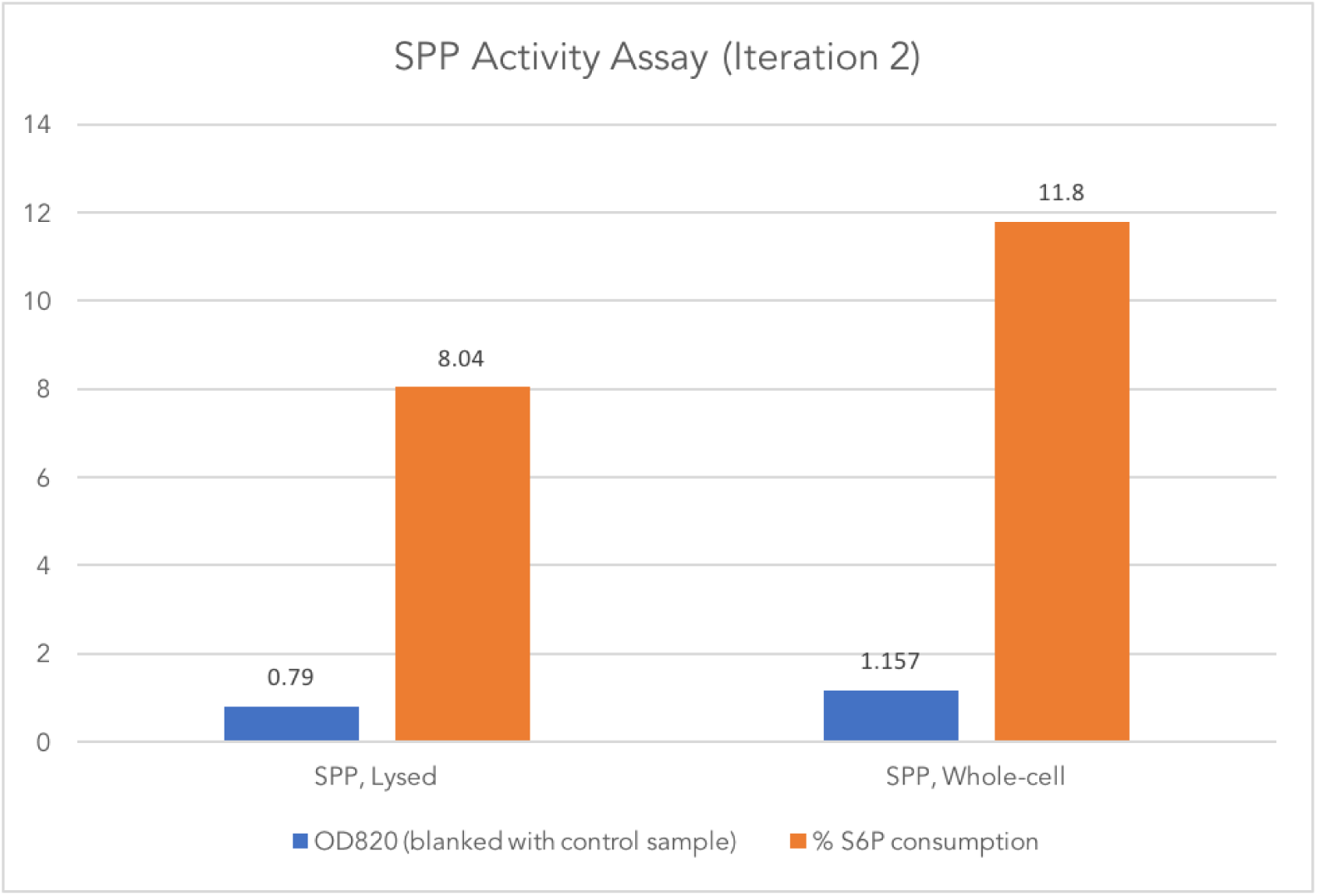
Ammonium Molybdate Assay with 8.04% (intact-cell) & 11.8% (lysed) S6P to Sucrose Conversion in SPP-transformed 1A976 *B. subtilis*

Following our “Agarose Microbead Encapsulation” protocol, we report an optimal spore density of 10^9 endospores/mL. Encapsulated sucrose concentrations varying from 0% to 20% were tested. With the objective to maximize sucrose concentration, in order to extend the material’s ability to sustain the bacteria upon germination, we report that the maximum sucrose concentration we were able to achieve was of 10%. At higher concentrations we observed microbead malformation and poor endospore encapsulation efficiency. Fig. 6 demonstrates the range of shapes and sizes of the synthesizes microbeads.

**Figure 6:**
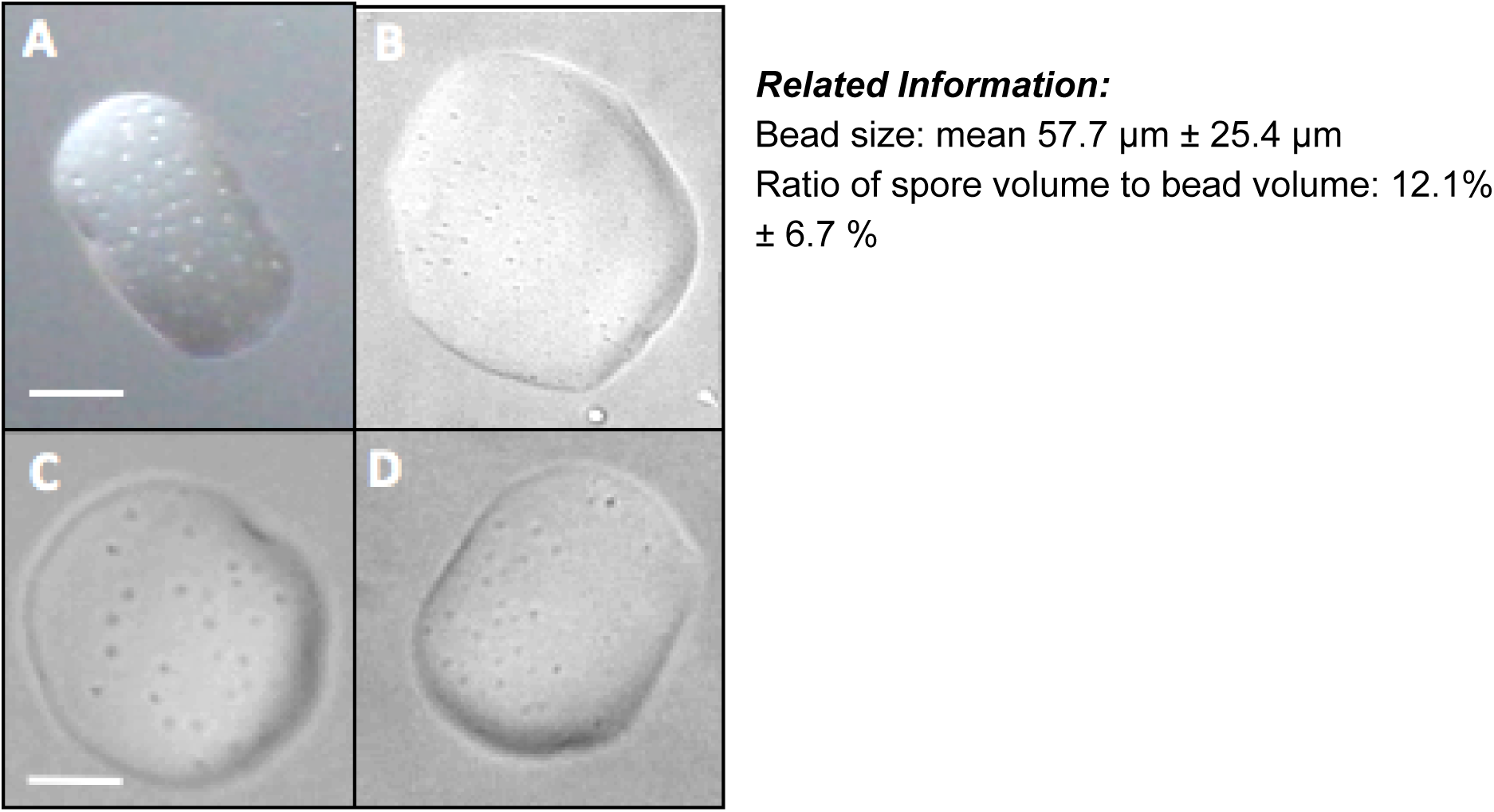
The images A-D were obtained by phase contrast microscopy. Shown, are *B. subtilis* spores encapsulated in low-gelling-temperature agar by gelation. A-B) Magnification: 630X, Scale bar: 20 µm C-D) Magnification: 1000X, Scale bar: 30 µm

To ensure the viability of the endospores post-encapsulation and post rupture, we used the Schaeffer-Fulton staining method to differentiate vegetative cells from endospores. We proceeded to encapsulate the stained endospores in agarose microbeads as described earlier. Subsequently, we simulated material rupture, by causing mechanical disintegration of the microbeads and inducing germination. Having observed the broken microbeads and their contents we report that the spores were viable and successfully underwent germination upon exposure to germinating conditions, as shown in Figure 7 below.

**Figure 7:**
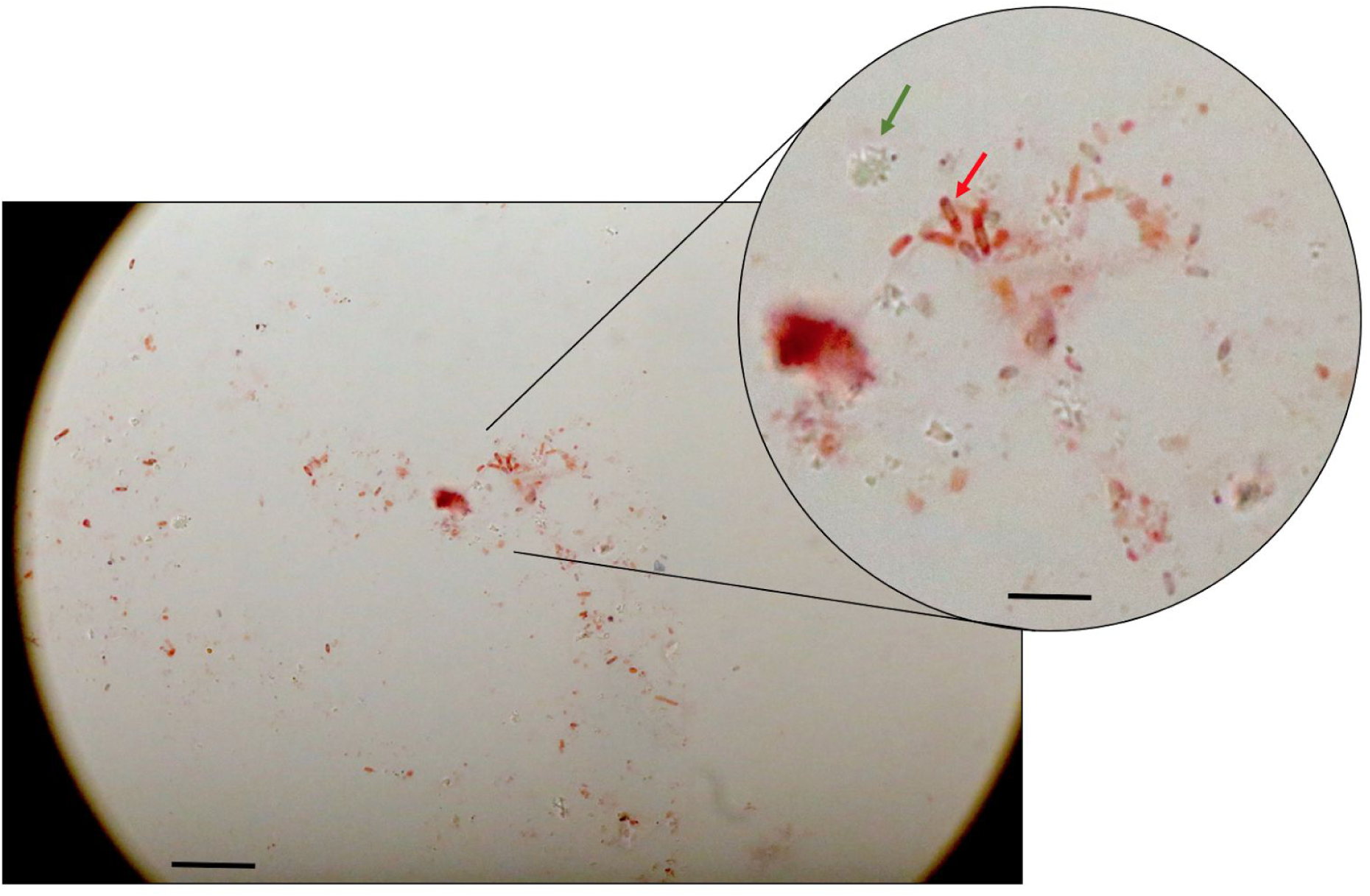
The image on the left was obtained as follows: Endospores of *B. subtilis* (strains 168, 168+trpC, PY79, SMY) were encapsulated in low-gelling-temperature agarose microbeads (mean 57.7 um ± 25.4 um) by gelation. The microbeads were crushed to release the embedded bacterial spores. The spores and potential vegetative cells were stained by the Schaeffer-Fulton differential staining method using malachite green and safranin. Mature spores stained green (green arrow), sporangia stained red (red arrow). Immature spores are visible within the sporangia. Broken pieces of agar microbeads are also visible in the sample (left on zoomed image). Scale bar on large image: 20 µm (1000X) Scale bar on zoomed image: 5 µm (4000X)

### BioBactery Results/Discussion

In order to assess the functionality of our proteorhodopsin within the voltage protein, we grew liquid cultures of NEB5α with and without our plasmid containing the voltage protein gene under the control of an arabinose inducible promoter. All cultures were grown in LB in the presence of 10µM all-trans retinal at 37°C in dark 15 mL Falcon tubes. The cultures containing our plasmid were grown with 0.2% arabinose to induce the production of our protein and with chloramphenicol, as our plasmid also harbored a resistance gene to this antibiotic.

On testing day, the OD600 of each culture was measured. Based on this measurement, a calculated volume of culture was spun down such that when resuspended in 10mL of the testing medium, the OD600 was 0.5. Once the cells were pelleted, the growth medium was discarded leaving only the pellet. Approximately 25 minutes before testing, pellets were resuspended in 10 mL of testing medium which was comprised of LB with sodium hydroxide added to adjust the pH to 9.04. We chose to work in an alkaline pH because the pH of the native ocean environment of SAR86 γ-proteobacteria is alkaline with a pH of ~8.2.^4^

For testing we illuminated each sample through 4 layers of green cellophane paper with a 2000W Xenon lamp with a UV filter, see Figure 2. Samples were exposed to alternating light and dark periods, 2 minutes each for a total of twelve minutes. pH measurements were recorded every 30 seconds with a pH meter accurate to 3 decimal places on a continuous read. Some samples were also exposed to varying concentrations of silver nitrate in an effort to stop cellular respiration which contributes to the proton motive force. If this proton motive force is already high, proteorhodopsin may be unable to pump protons according to Walter et al.^10^

Results of the testing show that the culture expressing the Voltage protein dropped pH significantly more than the control culture, see Figure 2. This demonstrates that the rhodopsin was pumping protons out of the *E. coli* in response to light. Interestingly, the addition of the respiration inhibitor, silver nitrate, did not seem to affect the pH drop. With the conclusion of this test, we confirmed that the rhodopsin domain is functional.

**Figure 2:**
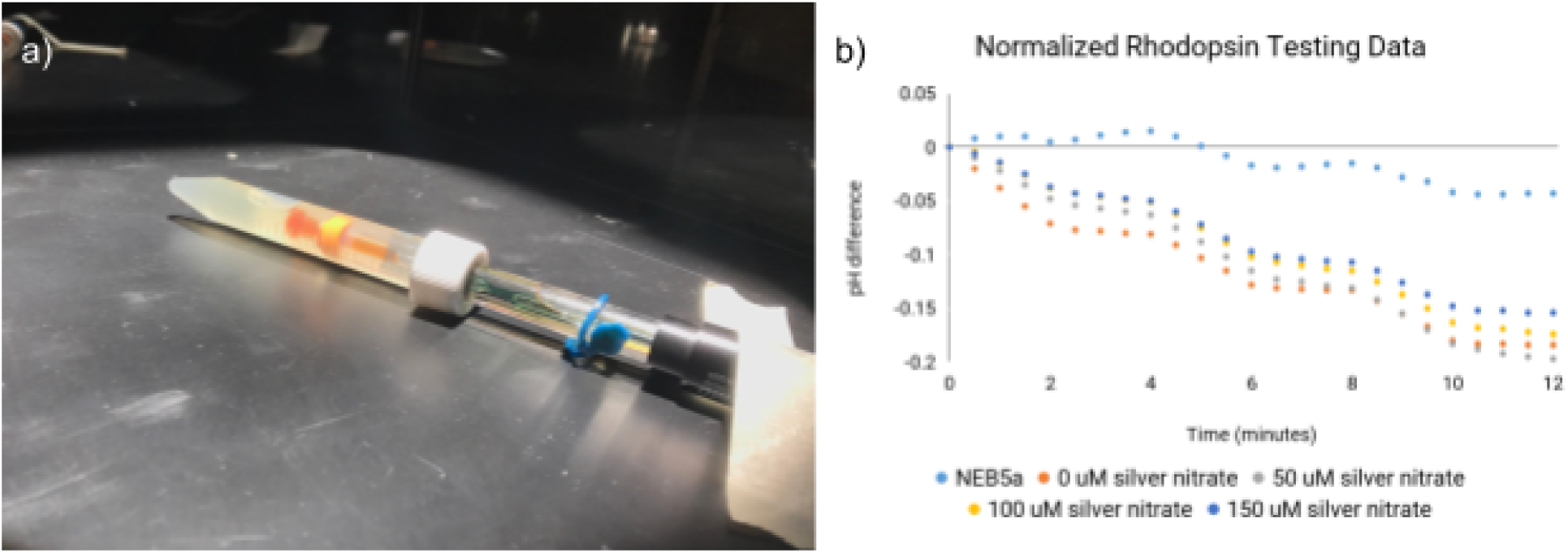
a) In our experimental setup, we inserted a pH probe into our bacterial cultures and exposed them to a 2000W Xenon lamp, two minutes on and two minutes off for twelve minutes. b)pH measurements were taken every 30 seconds to accuracies of three decimal places. Values were adjusted such that the data plotted is the pH difference from the start of the experiment.

To observe the localization of the Voltage protein, we observed the *E. coli* transformed with the plasmid containing the voltage protein with a fluorescence equipped microscope. After protein initial protein induction and observation, we were able to observe fluorescence; however, we were only able to observe slight polar localization at best, see Figure 3. We tried to vary the induction time, observing cultures after 30, 60, 120, and 180 minutes of induction by 0.2% arabinose, and we tried to vary the the amount of arabinose added from 0.01%, 0.1% and 0.2%, observing after 4 hours. Unfortunately, none of these attempts were successful in getting the voltage protein to completely unipolarly localize. In future efforts, it may be useful to try different linkers between the rhodopsin and the cPT region.

**Figure 4:**
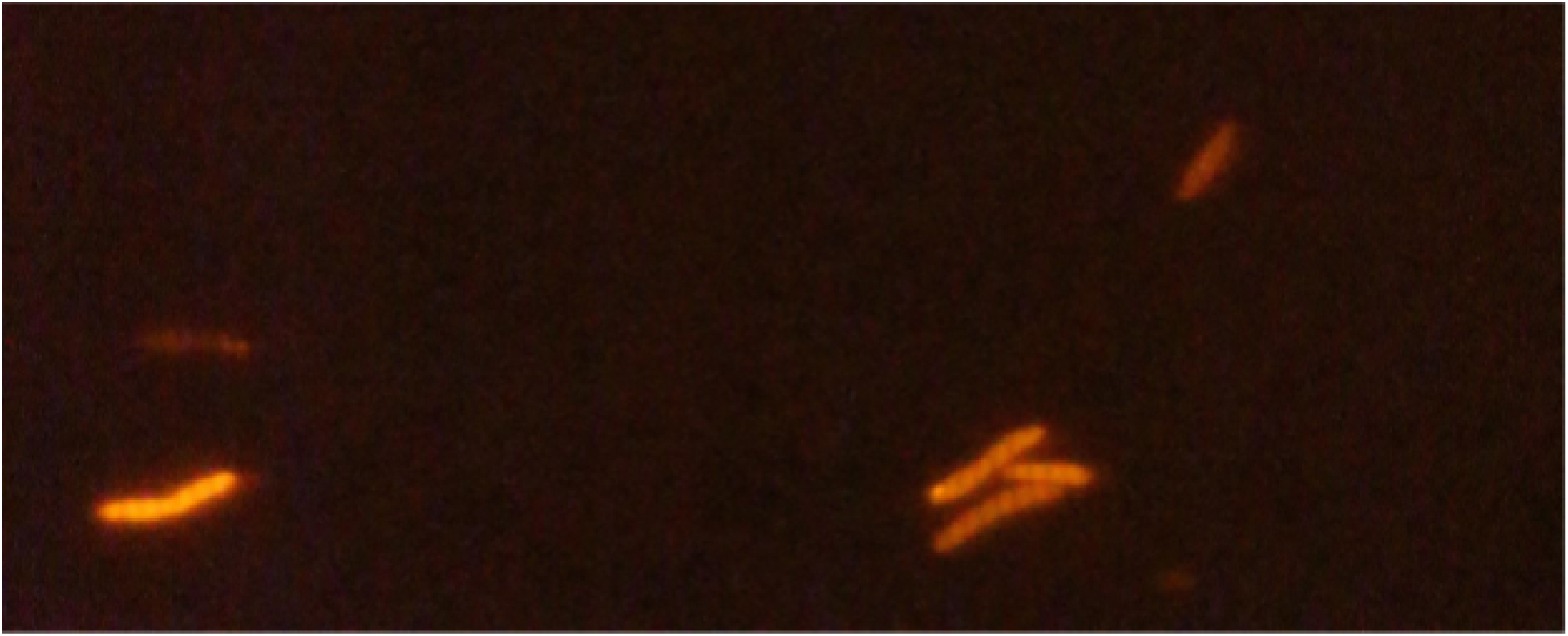
Image of voltage protein at 100x under fluorescent microscope, A slight unipolar focus forms in one *E. coli* cell.

The first method of orienting the *E. coli* we pursued was chemotaxis. At first we were discouraged because *E. coli* are peritrichous bacteria, meaning their cell surface is covered in flagella (Pratt et al.). This subsequently means that if we induced the cells to swim in one direction, there would be no guarantee with which side they would swim. Because our voltage construct theoretically localizes to the old pole of *E. coli*, having random directional swimming would mean that the ions would be pumped in a random direction relative to the microfluidic device. However, we then found a paper that found that 66% of the time, *E. coli* swim with their old poles behind them (Ping et al.). Although not completely polar, the skew of *E. coli*’s swimming pattern toward swimming with their old pole behind them was promising. We then conducted a chemotaxis experiment where we incubated *E. coli* strain W in M9 minimal media (“M9 Minimal Media Recipe”). The strain was selected for its flagella, as the normal lab strain NEB5a does not have flagella, and M9 minimal media was used because it has less nutrients than LB, meaning the *E. coli* would theoretically swim toward to the more nutrient-dense media.

**Figure 5:**
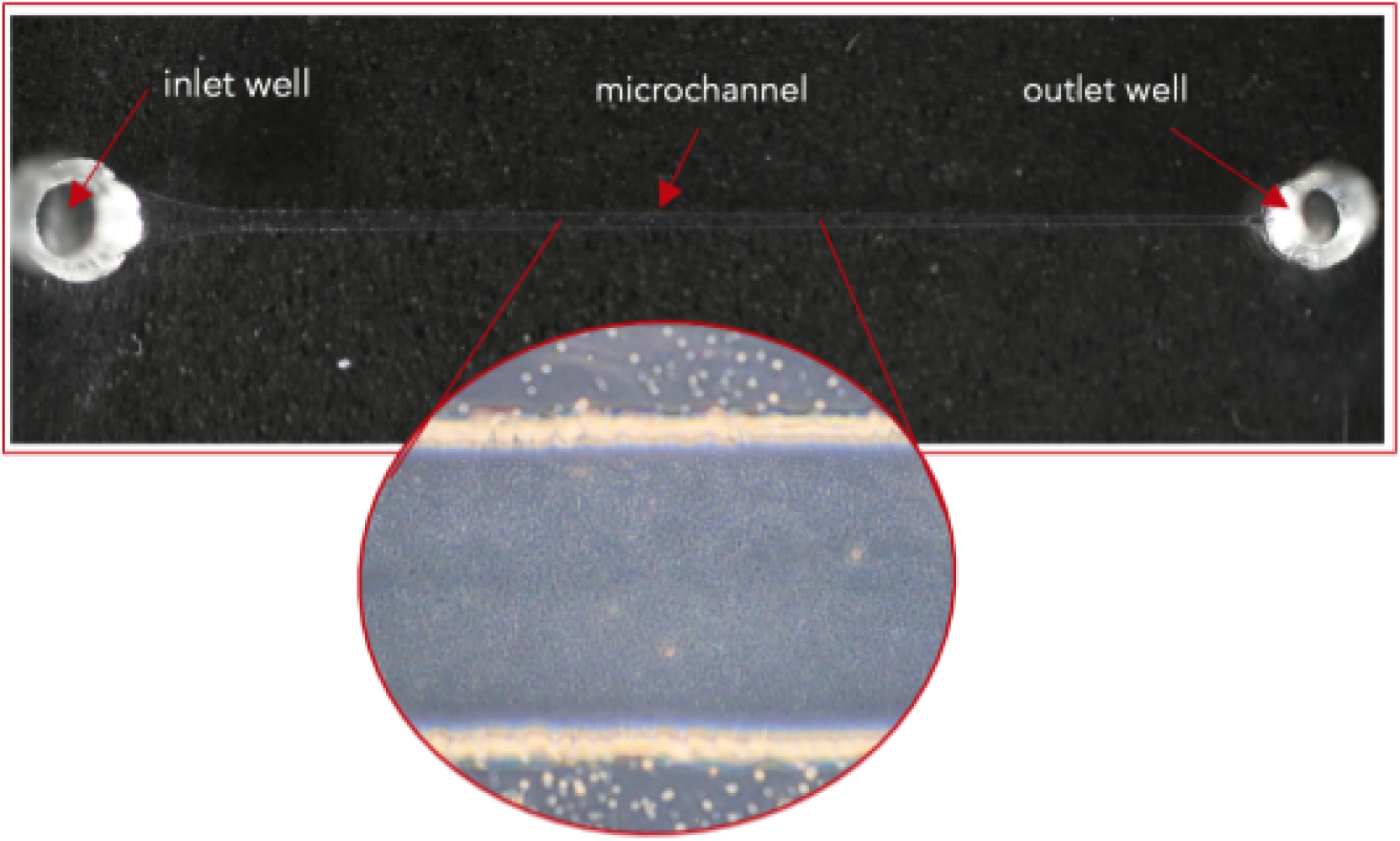
(a) image of microfluidic device, empty and then filled with *E. coli*. taken with the Zeiss microscope at 20X. (b) video of Chemotaxis experiment, filmed under the Zeiss microscope, wherein LB was used a chemoattractant in the outlet well and *E.* coli in M9 minimal media were placed in the inlet well at 20X. Random swimming patterns, rather than chemotactic swimming patterns, were observed.

We then pipetted approximately 15 μl of *E. coli* W in M9 minimal media into the inlet well of our microfluidic device, and 10 μl of LB, or lysogeny broth, into the outlet well. We then took the video in Figure 5. Unfortunately, we observed random motion of the *E. coli*, not directed movement toward the chemoattractant of LB. We therefore moved onto our second idea for orientation, mechanical orientation.

To test the polar localization of our biotinylated IcsA for mechanical orientation, we used a fluorescent streptavidin dye to bind to the outer membrane of our *E. coli*. We chose this method of verifying polar localization because fusing IcsA with a fluorescent protein causes it to localize intracellularly as opposed to extracellularly. With a generous gift from Thermo Fisher Scientific, we used Thermo Fisher Scientific’s Streptavidin, Alexa Fluor 405 conjugate to visualize the polar localization of IcsA (“Streptavidin, Alexa Fluor”). For this protocol, we tested strain W and NEB5α as a negative control. NEB5α was used as a negative control because it does not have a complete LPS, and therefore has too much membrane fluidity to see polar localization of IcsA. Strain W, therefore, was hypothesized to have polar localization of IcsA, and therefore fluorescence only at the old pole of the *E. coli*. After growing up the cultures overnight, the cultures were induced with IPTG at 0.2 mM for 5 hours. They were then spun down and resuspended in DPBS twice, and then incubated at 4 °C on a rotator for 30 minutes. The samples were then visualized at 20X and 40X under an Evos microscope. As expected, there was significantly more protein visualization and polar localization in strain W compared to NEB5a, as shown in Figure 6. For future experimentation to maximize the number of cells expressing IcsA and polar localization, the amount of fluorescent streptavidin dye could be varied, the level of IPTG induction could be manipulated, and the time given for the protein to fold could be studied.

**Figure 6:**
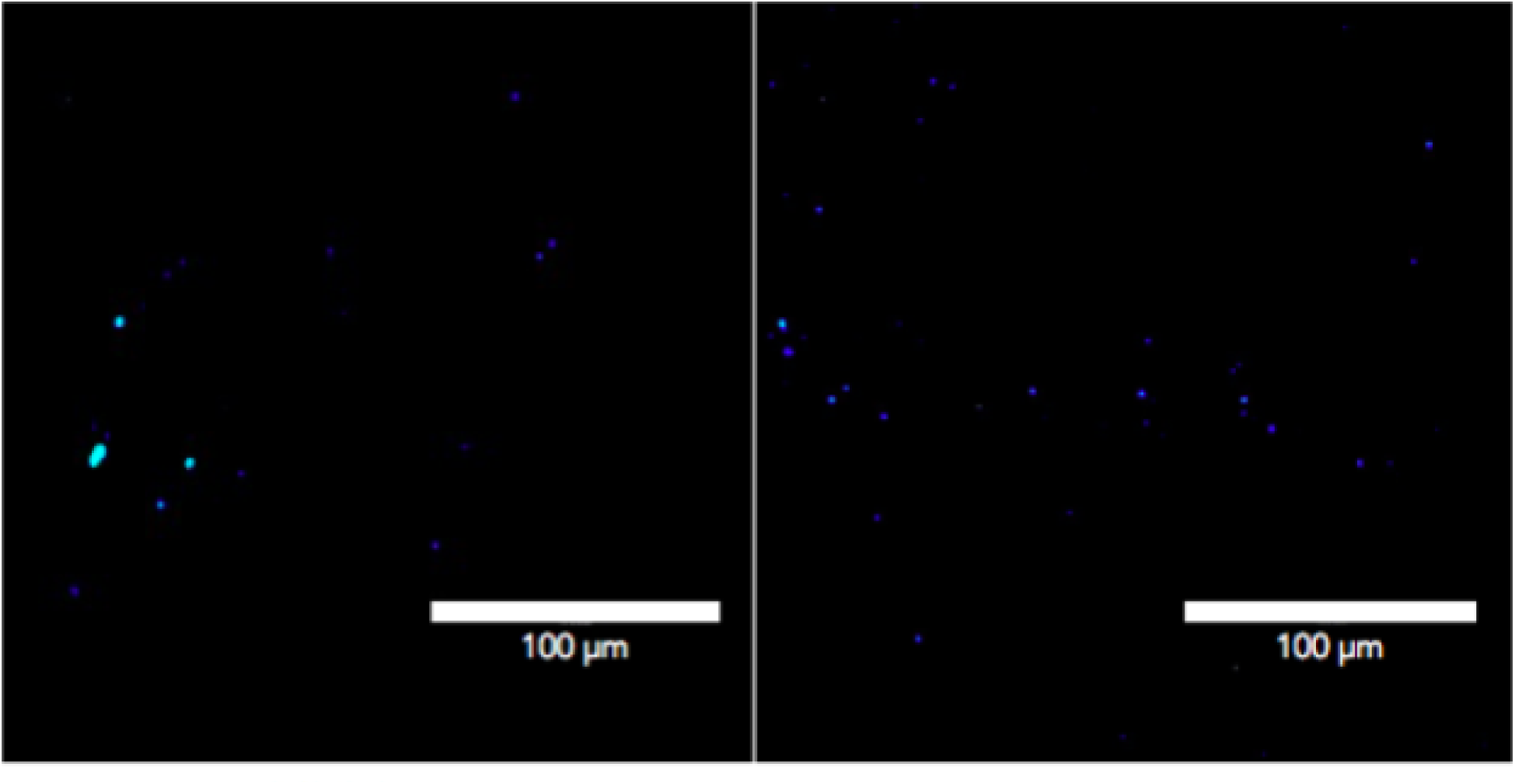
a) Shows fluorescent tagged-lcsA expressed in *E*. *coli* NEB5a, taken at 40X in an Evos digital microscope. b) Shows fluorescent tagged-lcsA expressed in *E. colí* strain W. Both images taken at 40X in an Evos digital microscope, with an 100 um scale bar.

The second assay we used to validate the functionality of our IcsA construct was magnetic bead attachment. The aim of this assay was to demonstrate that magnetic beads could be attached to our *E. coli*, and a positive result would be movement of the *E. coli* in response to an applied magnetic field. To do so, we purchased MojoSort streptavidin coated magnetic beads with a diameter of 30 nm that would be attached to our *E. coli* to orient them under a magnetic field (“MojoSort™ Streptavidin Nanobeads”). The small size of the magnetic beads was chosen specifically to counteract diffusion in the biobactery setup, as we wanted *E. coli* packed tightly together to promote unidirectional ion flow. We grew up liquid cultures of our IcsA-expressing *E. coli* overnight in both NEB and W strains, and measured the OD600 to determine the volume needed for 10^9^ cells. The cells were then washed twice with DPBS + 0.1% Tween 20, and then incubated with 5% or 10% of the magnetic bead solution to facilitate binding. After incubation on ice for 30 minutes, the cells with magnets were spun down and resuspended in the DPBS-Tween 20 solution. They were then placed in NEB magnetic separation rack for 5 minutes. At this step, we could see the magnetic beads and a small cell pellet separate out from solution closest to the magnet on the rack (Figure 7). The supernatant was taken out of the tube, and the magnet and cell pellet was then resuspended in the DPBS solution. Both the supernatant and the pellet were spun down and resuspended in a final volume of 100 μl of DPBS. Both solutions were prepared for microscopy. Because the beads are too small to visualize under the Evos microscope, we placed a strong magnet found in the teaching lab at the top of the slide to induce orientation of our *E. coli*, using their movement as the indicator for the magnetic bead attachment. We took the video in Figure 5 as evidence of the magnetic bead attachment, where we saw clear taxis of the *E. coli* toward the strong magnet. With these results, we have proven that we are able to attach magnetic beads to our *E. coli* via a biotin-streptavidin bond between IcsA and streptavidin covered magnetic beads.

**Figure 7:**
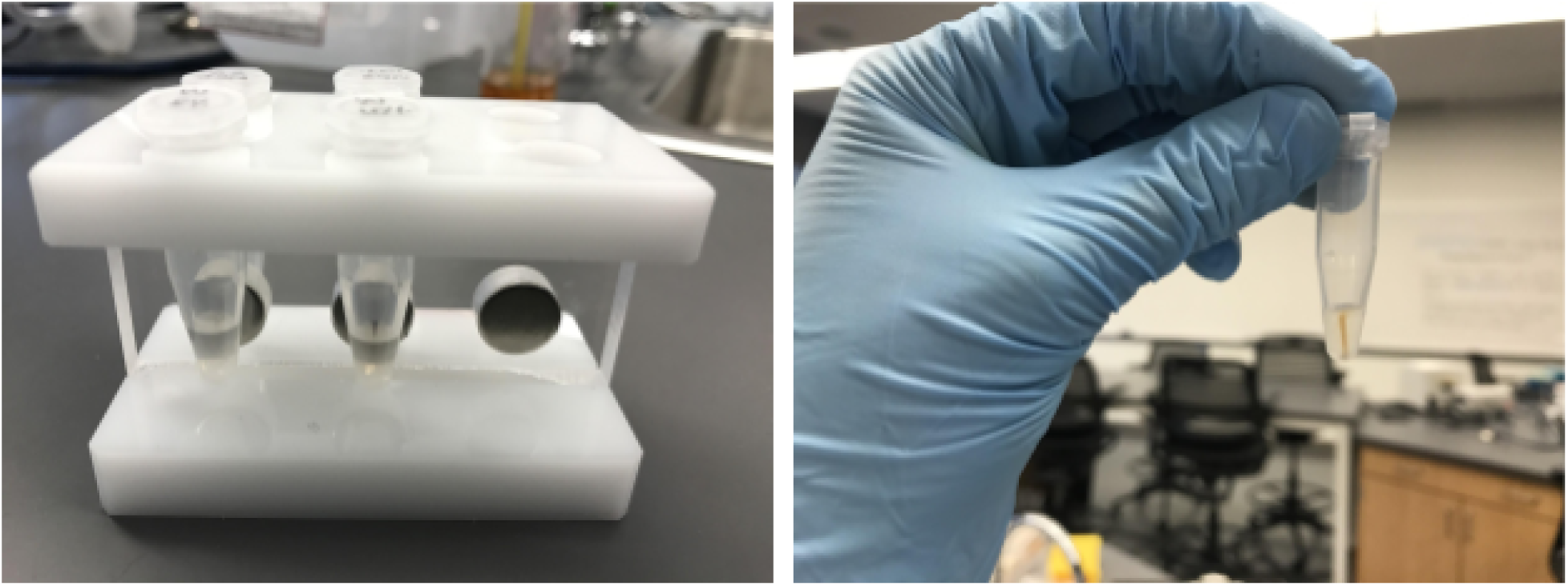
Visualization of magnetic bead attachment. a) Setup of samples in magnetic separation rack, where a pellet of magnetic beads conjugated to cells can be seen. b) One sample clearly showing the pellet of magnetic beads and cells. c) Video demonstrating the attachment of magnetic beads to IcsA-expressing *E. coli in* NEBSa strain. This sample was incubated with 10% of the Mojosort nanoscale magnetic bead solution, and visualized at 40X under the Evos microscope. A strong magnet was placed above the slide (out of frame), and the video demonstrates taxis toward the magnet 30 seconds after It has been placed above the slide.

Through the orientation component of this project, we have shown that we are able to orient *E. coli* in solution. We have yet to demonstrate this orientation in a microfluidic device, and this is a critical next step. Through the voltage component, we have show the functionality of our proton pump, but have achieved limited success in polarly localizing this protein. This may be due to protein misfolding with the specific linker we used, or it could be a fundamental limit in that the cPT region may not be able to unipolarly localize inner membrane proteins. In any case, further experimentation with the linker between proteorhodopsin and the cPT region would be a good next step.

Once we demonstrate orientation in a microfluidic device and achieve polar localization of the voltage protein, we will be ready to build our BioBactery. By transforming and expressing both constructs into our complete LPS *E. coli*, we would theoretically be able to generate a voltage drop across the length of a microfluidics channel. By attaching the magnetic beads to these cells, orienting them in the microfluidic device, and exposing them to green light to activate the proteorhodopsin, we should observe proton flow from one side of the channel to the other, thereby generating a voltage which could be used as a source of power. The power generated could potentially power small biosensors or one time use electronics at first, but with proper optimization, it could be scaled for a larger purpose. All in all, while we have yet to synthesize both aspects of this project, we have demonstrated successes in both components and are on our way to building a functional BioBactery.

## Acknowledgements

We’re very grateful for financial support from Stanford VPUE Grant for Undergraduate Research, Brown University UTRA, NASA Space Technology Mission Directorate Early Stage Innovation, the California Space Grant Consortium, and the Rhode Island Space Grant Consortium. Part of this work was performed at the Stanford Nano Shared Facilities (SNSF), supported by the National Science Foundation under award ECCS-1542152.

Special Thank You:

Dr. Marcia Goldberg at Harvard University - The team would like to acknowledge Dr. Goldberg’s instrumental guidance to facilitate our understanding of IcsA, for use in bioBactery. Additionally, we would like to thank her for sharing DNA samples.

Dr. David Shintani at University of Nevada, Reno - For his help and the gracious offer of use of his facilities for the molecular analysis of the synthetic latex generated in BioRubber.

Dr. Colleen McMahan at the United States Department of Agriculture - For her input on the enzymatic synthesis and degradation of latex.

Dr. Gary Wessel and Dr. Jim Head at Brown University for their input and advice.

Contributions: Conceptualization, Methodology, Writing - Original Draft Preparation
Contributions: Conceptualization, Formal Analysis, Methodology, Investigation, Writing - Original Draft Preparation
Contributions: Conceptualization, Data Curation, Investigation, Visualization, Writing - Original Draft Preparation, Writing - Review & Editing
Contributions: Conceptualization, Investigation, Methodology, Writing - Original Draft Preparation
Contributions: Conceptualization, Investigation, Methodology, Writing - Original Draft Preparation, Writing - Review & Editing
Contributions: Conceptualization, Investigation, Methodology, Writing - Original Draft Preparation
Contributions: Conceptualization, Data Curation, Formal Analysis, Investigation, Methodology, Visualization, Writing - Original Draft Preparation
Contributions: Conceptualization, Investigation, Methodology, Writing - Original Draft Preparation
Contributions: Conceptualization, Investigation, Supervision, Writing - Original Draft Preparation, Writing - Review & Editing
Contributions: Conceptualization, Methodology, Supervision, Writing - Review & Editing
Contributions: Conceptualization, Data Curation, Formal Analysis, Investigation, Methodology, Visualization, Writing - Original Draft Preparation
Contributions: Conceptualization, Methodology, Supervision, Writing - Review & Editing
Contributions: Supervision, Methodology, Writing - Review & Editing
Contributions: Supervision, Writing - Review & Editing
Contributions: Funding Acquisition, Resources, Methodology, Supervision
Contributions: Conceptualization, Funding Acquisition, Project Administration, Resources, Supervision, Writing - Review & Editing

